# Mechanistic Insights into the Inhibition of Recombinant Hepatitis E Virus Papain-like Cysteine Protease by *Khaya grandifoliola* Hydro-Ethanolic Extract: UHPLC-MS Profiling, Enzyme Kinetics, Computational Modeling, and Cell-Based Assays

**DOI:** 10.1101/2025.08.17.670703

**Authors:** Arnaud Fondjo Kouam, Cromwel Tepap Zemnou, Armel Jackson Seukep, Babayemi Olawale Oladejo, Brice Fredy Nemg Simo, Kerinyuy Juliene Kongnyuy, Armel Gaelle Kwesseu Fepa, Elisabeth Menkem Zeuko’o, Frédéric Nico Njayou, Paul Fewou Moundipa

## Abstract

**Ethnopharmacological Relevance:** Hepatitis E virus (HEV), a significant cause of liver diseases, lacks specific treatments. *Khaya grandifoliola* C.DC (Meliaceae), used in ethnomedicine to treat infections and liver-related ailments, shows promise as an antiviral agent. While its efficacy against hepatitis C virus (HCV) is known, its effects on HEV remain underexplored.

**Aims of the Study:** This study evaluates *K. grandifoliola*’s hydro-ethanolic extract (KHE) as a potential source of HEV inhibitor, focusing on the HEV papain-like cysteine protease (HEV-PLpro).

**Materials and Methods:** Phytochemical profiling of KHE was performed using a Waters-Acquity UHPLC-MS system. The recombinant HEV-PLpro, expressed in baculovirus-infected insect cells and purified via nickel-affinity and size-exclusion chromatography, was used to screen KHE’s antiviral activity. The IC_50_ and inhibition mechanism were determined using the fluorogenic substrate Z-RLRGG-AMC. Molecular docking and dynamics simulations predicted interactions and analyzed the stability of KHE’s compounds with HEV-ORF1 (PDB ID: 6NU9). Additionally, HEV replication inhibition and cytotoxicity were evaluated in Huh7.5 cells using a Gaussia luciferase reporter system and MTT assay, respectively.

**Results:** Eighteen compounds comprising flavonoid-O-glycosides, polyflavonoids, phenolic and terpene glycoside among other were successfully identified. KHE exhibited mixed-type inhibition of HEV-PLpro proteolytic activity, with an IC_50_ of 20.86 µg/mL, comparable to ribavirin (19.13 µg/mL), acting as competitive inhibitor. Notably, quercetin-3-[rhamnosyl-(1→2)-α-L-arabinopyranoside] and quercitrin exhibited stronger binding affinities (−7.68 and −7.43 kcal/mol) and greater structural stability, robustness, and compactness than ribavirin (−4.87 kcal/mol). In cell-based assays, KHE suppressed HEV replication more effectively than ribavirin (ICLL: 12.75 µg/mL vs. 17.96 µg/mL), with no cytotoxicity at ≤100 µg/mL.

**Conclusion:** Our results demonstrate that KHE exhibits potent anti-HEV activity by inhibiting both viral protease function and replication, likely attributable to its flavonoid-rich composition. Its mixed-type inhibition mechanism and favorable safety profile underscore its potential as promising source of lead candidate for HEV therapeutics. This study bridges traditional medicine and modern pharmacology, supporting further exploration of *K. grandifoliola* for antiviral drug development.

## 1. Introduction

Hepatitis E virus (HEV) remains a significant global public health concern, particularly in developing countries with inadequate sanitation infrastructure. As an emerging pathogen, HEV causes outbreaks of acute hepatitis and sporadic cases, placing substantial burdens on healthcare systems and affected populations (Raji et al., 2022; WHO, 2025). Annually, HEV is estimated to infect approximately 20 million people worldwide, resulting in nearly 60,000 deaths, with pregnant women and immunocompromised individuals at heightened risk (Nimgaonkar et al., 2018; WHO, 2025; Wu et al., 2020).

HEV is a non-enveloped, single-stranded, positive-sense RNA virus belonging to the *Hepeviridae* family (Wallace et al., 2019). Its approximately 7.2 kb genome contains three Open Reading Frames (ORFs). ORF2 encodes the viral capsid, while ORF3 encodes a multifunctional protein involved in viral morphogenesis and pathogenicity of new viral particles. ORF1 is translated into the nonstructural viral polyprotein featuring several functional proteins, including the methyltransferase enzyme, the Y domain, RNA-dependent RNA polymerase, and notably, the papain-like cysteine protease (HEV-PLpro), which is crucial for viral replication (Karpe, 2024; Kenney and Meng, 2019; Wang and Meng, 2021). The HEV-PLpro processes viral polyproteins, a step indispensable for the maturation and infectivity of new virions (Glitscher et al., 2024; LeDesma et al., 2019). Additionally, it interferes with host immune responses by blocking type I interferon induction, suggesting a role in immune system evasion (Kim and Myoung, 2018; Kim et al., 2016). Due to its essential role in the viral lifecycle, inhibiting HEV-PLpro activity presents a promising antiviral strategy.

Despite its health impact, there are no licensed vaccines or specific antiviral therapies for HEV infection. Current treatment options mainly involve ribavirin combined with pegylated interferon, which can cause side effects such as flu-like symptoms, fatigue, hematologic abnormalities, gastro-intestinal disorders, and neuropsychiatric issues (Davoodi et al., 2018; Sulkowski et al., 2011; Wu et al., 2016). Accordingly, the health risks posed by HEV infection, coupled with the limited treatment options, prompt the urgent need for novel antivirals, particularly from natural sources like medicinal plants. Nature offers an abundant repository of bioactive compounds, many of which have historically served as templates for drug discovery (Beutler, 2009; Vaou et al., 2025). Plants constitute a rich source of phytochemicals with diverse biological activities, including antiviral, antioxidant, anti-inflammatory, and immunomodulatory effects (Behl et al., 2021; Lin et al., 2014). Exploring plant extracts and their constituents has gained interest as a complementary approach to antiviral therapy, especially since ethnobotanical practices may harbor compounds capable of modulating viral enzymes or interfering with viral replication (Adeosun and Loots, 2024; Rodríguez-Negrete et al., 2024).

One such promising botanical resource is *Khaya grandifoliola* C.DC, commonly known as African mahogany, which belongs to the *Meliaceae* family. Widely distributed across West Africa, it has been traditionally used to treat infections, inflammations, fever, malaria, and liver-related ailments (Danquah et al., 2019; Njayou et al., 2016). Phytochemical investigations of *Khaya grandifoliola* have revealed a complex mixture of bioactive constituents, including limonoids, triterpenoids, flavonoids, and other phenolic compounds (Kouam et al., 2017). These phytochemicals have demonstrated a range of pharmacological properties, such as antimicrobial, anti-inflammatory, antioxidant, antiviral, and antihepatotoxic activities (Kouam et al., 2022, 2024b, 2019; Mukaila et al., 2021). Interestingly, extracts and limonoids from *K. grandifolila* have demonstrated inhibitory effects against hepatitis C virus (Galani et al., 2016; Kouam et al., 2021). Despite these promising biological activities, little is known about the effect of *K. grandifoliola* on HEV.

Advances in analytical chemistry and computational biology have paved the way for a more systematic understanding of plant phytochemicals and their mechanisms of action (Khoo et al., 2018; Seukep et al., 2025; Zemnou, 2025). Techniques such as ultra-high-performance liquid chromatography coupled with mass spectrometry (UHPLC-MS) enable detailed phytochemical profiling, helping identify and quantify bioactive compounds, potentially responsible for the observed biological activities (Atanassova et al., 2025). Concurrently, enzyme kinetics studies provide insights into how these compounds interact with viral enzymes, revealing potential modes of inhibition (Kouam et al., 2024a, 2025a). Complementary computational studies, including molecular docking and molecular dynamics simulations, further delineate the binding interactions at the molecular level, informing structure-activity relationships and guiding the design of more potent inhibitors (Jain et al., 2025; Zemnou, 2025).

Given the importance of developing effective HEV antiviral agents, this study explored the inhibitory potential of the hydro-ethanolic stem bark extract of *K. grandifoliola* on HEV-protease activity. The specific aims being to characterize the phytochemical composition of the extract through UHPLC-MS, evaluate its inhibitory effects on the viral protease through enzyme kinetics, and explore the underlying mechanisms via computational modeling, and evaluate the *in vitro* antiviral potential.

## 2. Materials and Methods

### 2.1 Cells, viruses and culture conditions

Insect cells: *Spodoptera frugiperda* Sf9 cells line (Invitrogen, CA, USA) were used for the production and amplification of the recombinant baculovirus using the pFastBac^TM^1 plasmid vector (Invitrogen, CA, USA), and *Trichoplusia ni* High Five cells line were used for large scale production of the recombinant protein. They were maintained in Insect-XPRESS Protein-free Insect Cell Medium (Cat N° 12-730Q, LONZA, Verviers, Belgium) supplemented with penicillin (100 UI/mL) and streptomycin (100 µg/mL) in suspension culture at 27°C under shaking at 150 rpm. Huh7.5 cells line, used for the in vitro antiviral assay, were grown Dulbecco’s Modified Eagle Medium (DMEM) (Gibco, Carlsbad, CA, USA) containing 10% Fetal Bovine Serum (FBS), L-Glutamine (2 mM), penicillin (100 UI/mL), streptomycin (100 µg/mL), and Fungizone (2.5 µg/mL) in humidified atmosphere of 5% CO2 at 37°C. pJE03-1760F/P10-GLuc plasmid, encoding the viral genome of a chimeric strain of HEV genotype 3 (Nishiyama et al., 2019b), was kindly gifted by the Institute of Microbiology, Chinese Academy of Sciences (Beijing, China). It contains a luciferase gene of Gaussia princeps, inserted between the ORF2 and ORF3 genes. In transfected cells, pJE03-1760F/P10-GLuc has been demonstrated as suitable reporter virus assay to evaluate the inhibitory potential of drug candidates targeting HEV replication, by measuring the bioluminescence of GLuc activity secreted in the incubation medium (Nishiyama et al., 2019a).

### 2.2 Collection of plant material, authentication and preparation of extract

The studied plant material comprised the stem bark of *K. grandifoliola*, which was collected in March 2023 from the Foumban locality in the Noun Sub-Division of the West Region of Cameroon. The plant’s nomenclature was verified using the online resource http://www.theplantlist.org, and the botanical identification was conducted at the Cameroon National Herbarium, where a voucher specimen was preserved under the reference number HNC/23434 YA. The extraction was performed as previously described (Kouam et al., 2024b). Briefly, the collected plant material underwent a washing process, followed by air drying and grinding. Subsequently, 100 grams of the resultant powder was subjected to maceration at ambient temperature (25L) with 1 liter of a solvent system composed of ethanol and water in a 70:30 (v/v) ratio for a duration of 24 hours. The mixture was then filtered using Whatman No.1 filter paper, and the residual material was re-extracted once. The filtrates were subjected to rotary evaporation at 65°C under reduced pressure to eliminate the ethanol content. The remaining extract was dried at 40L, yielding 21 grams of extract, which has been designated as KHE (hydroethanolic extract of *K. grandifoliola*).

### 2.3 Ultra-High Performance Liquid Chromatography-Mass spectrophotometry (UHPLC-MS) profiling of KHE

Ultra-High Performance Liquid Chromatography-Mass Spectrometry (UHPLC-MS) was used to determine the phytochemical composition of KHE, employing a Waters Acquity system coupled with a mass spectrometer, as previously described (Kouam et al., 2025b). Briefly, the instrument comprised a vacuum degasser, quaternary pump, auto-sampler, and photodiode array (PDA) detector, enabling simultaneous chromatographic and MS spectral data acquisition. Data were processed using Waters MassLynx 4.1 software. Electrospray ionization was performed in negative mode (15 V), scanning m/z 150–1500. A Waters HSS T3 column (2.1 mm × 100 mm, 1.7 µm particle size) was used for the separation of compounds, with a mobile phase of 0.1% formic acid in water (A) and acetonitrile (B), flowing at 0.3 mL/min under the following gradient conditions: 0–1 min (100% A), 1–9 min (28% B), 9–10 min (40% B), and 10–12 min (100% B). The column was kept at 55°C, and 2 µL of KHE at 1 mg/mL in methanol was injected. Identification of chemical compounds was done using MSDIAL and MSFINDER (RIKEN Center for Sustainable Resource Science: Metabolome Informatics Research Team, Kanagawa, Japan) by matching their respective m/z values, SMILES formulas and InChIKey (International Chemical Identifier) identifiers (Lai et al., 2018; Tsugawa et al., 2015). Supplementary file S1 provides additional details.

### 2.4 Production of recombinant HEV papain-like protease

#### 2.4.1 Construction of the expression plasmid vector

The coding sequence for the HEV papain-like protease (HEV-PLpro) domain (nucleotides 1319– 1801 of HEV genotype 2, accession number AF444002) was obtained from the NCBI GenBank database. This sequence was synthesized and codon-optimized for expression in the Bac-to-Bac baculovirus system in insect cells. The synthesized gene was amplified via PCR and cloned into the EcoRI and KpnI restriction sites of the pFastBac™1 vector. The expression construct included an N-terminal GP67 signal peptide to facilitate secretion of the recombinant protein into the culture medium and a C-terminal hexahistidine (6×His) tag for purification via nickel affinity chromatography. The expression plasmid vector was assembled by SynbioTechnologies (Beijing, China).

#### 2.4.2 Cloning of HEV-PLpro into Bac-to-Bac system and isolation of recombinant Bacmid DNA

The HEV-PLpro expression vector (pFastBac1-N-GP67-HEV-PLpro-C-6His) was transformed into competent *Escherichia coli* strain Max Efficiency™ DHα10Bac™ (Invitrogen), which contains the baculovirus shuttle vector (Bacmid). Then, 900 µL of Luria-Bertani (LB) medium was added to the transformed bacteria and incubated at 37°C while shaking at 250 rpm. After four hours, serial tenfold dilutions (10L¹, 10L², 10L³) were prepared, and 100 µL of each dilution were plated on LB agar supplemented with kanamycin (50 µg/mL), gentamicin (7 µg/mL), tetracycline (10 µg/mL), X-gal (100 µg/mL), and isopropyl-D-thiogalactoside (IPTG) (40 µg/mL) to select for positive clones (white colonies) over negative clones (blue colonies) (Fig. 2A), after 48 h of incubation at 37°C. To confirm the positive colonies, PCR amplification was performed using M13 primers (Forward: 5′-GTTTTCCCAGTCACGAC-3′; Reverse: 5′-CAGGAAACAGCTATGAC-3′), followed by agarose gel electrophoresis. Clones exhibiting a specific band at approximately 5000 bp (insert plus vector) (Fig. 2B) were selected for Bacmid DNA isolation. A single positive white colony was inoculated into 5 mL LB containing the antibiotics and incubated overnight at 37°C with shaking. Bacteria were pelleted by centrifugation (8000×g, 5 min), and Bacmid DNA was extracted using the TianPrep Mini-Plasmid Extraction kit (TIANGEN BIOTECH, Beijing, China) according to the manufacturer’s instructions.

#### 2.4.3 Generation and amplification of recombinant Baculovirus containing HEV-PLpro

Purified Bacmid DNA was transfected into Sf9 insect cells using FuGene 6 transfection reagent (Promega, Madison, USA). Briefly, Sf9 cells (approximately 0.8×10L cells/well at exponential growth phase) were plated in 6-well plates and allowed to attach for 15 minutes at room temperature. For each well, the transfection mixture contained 2 µg of Bacmid DNA, 16 µL of FuGene 6, and 200 µL of Insect-XPRESS medium. The mixture was vortexed and added to the cells after a 5-minute incubation at room temperature. After 6 hours at 27°C, the medium was replaced with fresh insect medium containing antibiotics. Cells were incubated and monitored daily until cytopathic effects (CPE) >75% were observed (typically after 48–72 hours). The supernatant, containing passage 1 (P1) virus stock with a low viral titer, was collected. P1 virus was amplified by adding 1.5 mL of P1 stock to about 10L Sf9 cells cultured in a 10 cm dish to produce P2 stock virus after 72 hours incubation at 27°C. Subsequently, high-titer P3 virus were generated by infecting approximately 10L cells with 1 mL of P2. For large-scale infections, P3 was further expanded to P4 virus (high viral titer used to infect High Five cells for protein expression) by infecting 450 mL of Sf9 cell suspension (density ≈ 1.9×10L cells/mL) with 3 mL of P3 virus.

#### 2.4.4 Expression and purification of recombinant HEV-PLpro

Large-scale expression of recombinant HEV-PLpro was performed using High Five cells. Briefly, four 2-liter flasks containing 400 mL of cell suspension at a density of 0.8 × 10^6^ cells/mL were incubated at 27°C under constant shaking at 150 rpm until the cell density reached approximately 1.9 × 10^6^ cells/mL. Subsequently, 100 mL of P4 virus was added to each flask, and the infected High Five cells were incubated for 60 hours. Afterward, the cells were collected and clarified by centrifugation at 8000 × g for 60 minutes at 4°C. The supernatant, which contained soluble recombinant HEV-PLpro, was filtered through a 0.22 µm filter membrane and applied to a HisTrap™ nickel-affinity column chromatography (GE Healthcare) pre-equilibrated with the binding buffer (20 mM Tris-HCl, 150 mM NaCl, 1 mM TCEP, pH 8.0). The target protein was subsequently eluted using a 30% elution buffer (20 mM Tris-HCl, 150 mM NaCl, 1 mM TCEP, 1000 mM imidazole, pH 8.0) with an AKTA Pure Purification System (GE Healthcare, Chicago, Illinois, USA). The recombinant HEV-PLpro was further purified by gel filtration chromatography using a HiLoad™ 16/60 Superdex 200 column (GE Healthcare) with gel filtration buffer (20 mM Tris-HCl, 150 mM NaCl, 2 mM DTT, pH 8.0). The purity of the collected fractions was assessed by sodium dodecyl sulfate polyacrylamide gel electrophoresis (SDS-PAGE), and fractions with high purity were selected, pooled, concentrated using a 10 kDa cut-off membrane (Amicon, Darmstadt, Germany), flash-frozen in liquid nitrogen, and stored at −80°C until use.

### 2.5 Assessment of the inhibitory effect of KHE on the proteolytic activity of the purified HEV-PLpro

#### 2.5.1 Determination of the kinetic parameters of the purified HEV-PLpro

The proteolytic activity of purified HEV-PLpro was evaluated using the fluorogenic substrate Cbz-Arg-Leu-Arg-Gly-Gly-7-amino-4-methyl coumarin (Z-RLRGG-AMC) (Bachem Biosciences, Bubendorf, Switzerland). The assay was carried out in a total volume of 100 µL in 96-well black plates. The reaction mixture comprised 80 µL of assay buffer (20 mM Tris-HCl, 150 mM NaCl, 2 mM DTT, pH 7.4), 10 µL of recombinant HEV-PLpro (final concentration 2 µM), and 10 µL of Z-RLRGG-AMC substrate at various concentrations (6.25, 12.5, 25, 50, 100, and 200 µM), diluted in assay buffer. The enzymatic activity was monitored at 35°C over 20 minutes by measuring fluorescence, which increased as AMC was released upon substrate hydrolysis. Fluorescence was recorded at an excitation wavelength of 350±20 nm and emission at 450±20 nm, with measurements taken every 80 seconds using a BMG LabTech ClarioStar multi-label plate reader (Ortenberg, Germany) with accompanying ClarioStar software (v6.20). For each substrate concentration, the initial velocity was calculated from the linear portion of the fluorescence versus time curve, expressed as Relative Fluorescence Units per minute (RFU·minL¹). A calibration curve, obtained by measuring fluorescence of free AMC at known concentrations (0.005 to 2.8 µM), was used to convert fluorescence units to the amount of AMC released. The initial velocities were finally expressed as the rate of AMC hydrolyzed (µM·minL¹). Kinetic parameters including the Maximum reaction velocity (Vmax) and Michaelis-Menten Constant (Km) were obtained by plotting Michaelis-Menten and Lineweaver-Burk graphs using GraphPad Prism (v8.0.2).

#### 2.5.2 Evaluation of KHE’s inhibitory effect on HEV-PLpro activity

The inhibitory effect of KHE on HEV-PLpro was assessed as previously described (Kouam et al., 2025a). Briefly, 70 µL of assay buffer, 10 µL of HEV-PLpro (2 µM), and 10 µL of KHE or ribavirin (used as a reference inhibitor) dissolved in 10% DMSO at various concentrations (0.01–100 µg/mL) were mixed and incubated at 35°C for 15 minutes. The reaction was initiated by adding 10 µL of Z-RLRGG-AMC (50 µM final concentration), and fluorescence was monitored for 20 minutes. The initial reaction rate for each inhibitor concentration was determined, and the inhibition percentage was calculated using the formula (1).

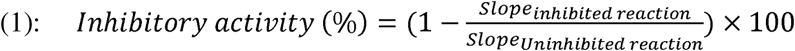

where “Slope” refers to the slope of the fluorescence vs. time curve. The inhibitory activity was plotted against inhibitor concentration, and the IC_50_ value (the concentration required to inhibit 50% of enzyme activity) was obtained via non-linear regression analysis (log [inhibitor] vs. response – Variable slope - four parameters) using GraphPad Prism.

#### 2.5.3 Determination of the mechanism of KHE on HEV-PLpro activity

To determine the type of inhibition exerted by KHE or ribavirin on HEV-PLpro, assays were performed with varying substrate concentrations (6.25, 12.5, 25, 50, 100, and 200 µM) and two inhibitor concentrations (½ and ¼ IC_50_). For each substrate concentration, the initial reaction rates with and without inhibitors were measured. Results were plotted as initial velocity versus substrate concentration, and enzyme kinetic analysis was performed using GraphPad Prism to identify the best-fit inhibition mechanism (competitive, uncompetitive, non-competitive, or mixed). Finally, secondary plots of 1/Vmax’ or Km’ against inhibitor concentration were used to determine the inhibitory constant (Ki) for each inhibitor.

### 2.6 Computational studies of identified compounds of KHE

#### 2.6.1 Molecular docking analysis of interactions between KHE compounds and HEV_ORF1

The crystal structure of the HEV_ORF1 non-structural protein, which includes the HEV-PLpro domain, was obtained from the Protein Data Bank (PDB ID: 6NU9). The binding site of ORF1, crucial for molecular docking studies, was determined by analyzing protein-ligand interactions within the PDB structure. The ligands used in this study were sourced from the PubChem database (https://pubchem.ncbi.nlm.nih.gov/) using their respective compound IDs. Before docking, the three-dimensional structures of all ligands were energy-minimized using Avogadro (https://avogadro.cc/) to ensure proper geometry. Molecular docking analysis was conducted using AutoDock 4.0 (http://autodock.scripps.edu/), a well-established software for predicting protein-ligand binding affinities (Zemnou, 2025). The Genetic Algorithm (GA) was employed with the following parameters: a population size of 300, 27,000 generations, 1,000,000 energy evaluations, and 100 independent GA runs to enhance docking accuracy. After docking the ORF1 protein with the respective compounds, protein-ligand interactions were analyzed and visualized using Discovery Studio (https://www.3ds.com/products-services/biovia/) to identify key binding residues and interaction patterns.

#### 2.6.2 Molecular Dynamic Simulation (MDS) of Lead Ligands from KHE

The best protein-ligand complexes from docking were further examined through Molecular Dynamics Simulation (MDS) to study the dynamic behavior of their atoms and stability. MDS was conducted using GROMACS-2023 (Abraham et al., 2015), with the Visual Dynamics interface (Vieira et al., 2023) supporting script creation. Ligand topologies were generated via ACPYPE (Kagami et al., 2023), utilizing the Amber99 force field (Wang et al., 2004) and Partial Charges based on ANTECHAMBER (Wang et al., 2006). Electrostatic interactions were addressed with the particle mesh Ewald (PME) method, employing a cutoff distance of 12 Å. Water molecules modeled with TIP3P (Jorgensen et al., 1983) were added to the simulation box. The system underwent a two-phase energy minimization: 2000 steps each of steepest descent and conjugate gradient until the force dropped below 1000 kJ·molL¹·nmL¹. Initial atomic velocities were assigned at 300 K using the Maxwell-Boltzmann distribution. Subsequently, the system was equilibrated with two 200 ps simulations under NVT and NPT ensembles. All complexes were then simulated freely at 300 K in the Gibbs ensemble with isotropic coupling at 1 atm pressure, with a 2 fs timestep. Hydrogen bonds and other chemical constraints were maintained using the SHAKE algorithm (Kräutler et al., 2001). During the simulation, parameters such as secondary structure content, radius of gyration (Rg), solvent-accessible surface area (SASA), intramolecular hydrogen bonds (H-bond), root-mean-square deviation (RMSD), and root-mean-square fluctuation (RMSF) were measured over 100 ns (Supplementary file S2).

### 2.7 Determination of the *in vitro* antiviral activity of KHE on HEV replication

#### 2.7.1 Linearization and in vitro transcription pJE03-1760F/P10-GLuc plasmid

Approximately 20 µg of the pJE03-1760F/P10-GLuc plasmid was linearized utilizing the FastDigest BamHI restriction enzyme (Catalog No. FD0054, Thermo Scientific) and subsequently purified through a standard phenol/chloroform extraction procedure. A 1 µg aliquot of the linearized DNA plasmid template was then transcribed into RNA employing the mMESSAGE mMACHINE™ T7 transcription kit (Catalog No. AM1344, Thermo Scientific). Upon completion of the transcription process, Turbo DNase (Catalog No. AM2238, Thermo Scientific) was utilized to degrade the DNA template, and the resultant RNA was purified through phenol/chloroform extraction followed by isopropanol precipitation. The concentration and purity of the isolated RNA were evaluated by measuring absorbance at 230, 260, and 280 nm using a NanoDrop spectrophotometer (ND-2000, Thermo Scientific).

#### 2.7.2 Transfection and bioluminescence-based antiviral assay

Huh7.5 cells were plated in 96-well plates at a density of 6.4 × 10^3^ cells per well in 100 µL of DMEM medium and incubated at 37°C. After 24 hours, the cells were washed with fresh medium, and transfection was performed by incubating the cells with 100 µL per well of the transfection complex. This complex contained 0.5 µg of pJE03-1760F/P10-GLuc RNA transcripts and 1 µL of Lipofectamine™ 2000 transfection reagent (catalog number 11668019, Thermo Scientific), which was diluted in serum-free OPTI-MEM medium (catalog number 31985062, Thermo Scientific). After 4 hours of incubation, the supernatants from both transfected and non-transfected (control) cells were removed, and the cells were washed twice with DMEM. They were then incubated with or without KHE or Ribavirin at final concentrations of 0.01, 0.1, 1, 10, 25, 50, and 100 µg/mL. After 72 hours, the cell culture supernatants were collected for the measurement of bioluminescence resulting from the secreted Gaussia luciferase.

#### 2.7.3 Measurement of *Gaussia* luciferase activity

GLuc activity was measured as previously reported (Kouam et al., 2021). Briefly, the cell culture supernatant from both transfected and non-transfected cells was clarified by centrifugation at 12,000 × g for 10 minutes at 25°C. The supernatant was then mixed with 0.25 volumes of Renilla 5X lysis buffer (Promega, Madison, USA). Bioluminescence was quantified in terms of Relative Light Units (RLU) utilizing the GLOMAX™ 20/20 Luminometer Tube Reader. This procedure entailed the mixing of 20 µL of the clarified cell supernatant with 50 µL of Gaussia luciferase assay reagent (New England BioLabs), followed by a 10-second integration period. For each concentration of inhibitors tested, the corresponding antiviral activity (% inhibition) was determined using formula (2):

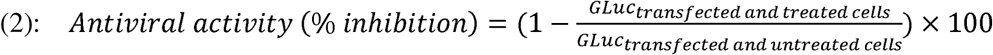

The IC_50_, defined as the concentration of an inhibitor that reduces GLuc bioluminescence values by 50% compared to transfected and untreated cells, was determined through non-linear regression analysis (Log [inhibitor] vs. response – Variable slope – four parameters) of the graph depicting the percentage of inhibition versus the concentration of inhibitors, utilizing GraphPad Prism software.

#### 2.7.4 Assessment of the cytotoxic effect of KHE on Huh7.5 cells

Huh7.5 cells were seeded in a 96-well plate at a density of 6.4 × 10^3^ cells per well and incubated for 24 hours at 37°C. Subsequently, the medium was replaced with fresh medium containing KHE or Ribavirin at various concentrations (0.1, 1, 10, 25, 50, and 100 µg/mL). After 72 hours of incubation, cell viability was assessed using the 3-(4,5-dimethylthiazol-2-yl)-2,5-diphenyl-2H-tetrazolium bromide (MTT) assay, as previously reported (Kouam et al., 2020). Cell viability was expressed as a percentage relative to the control group (untreated cells), with the viability of control cells considered to be 100%.

### 2.8 Statistical analysis

The results are expressed as means ± standard deviations derived from three independent experiments; each conducted in triplicate. To compare the IC_50_ values between the reference inhibitor (Ribavirin) and KHE, Student’s *t*-test was employed. Additionally, one-way analysis of variance (ANOVA) was utilized to assess the mean values across different treatment groups, followed by Bonferroni’s post hoc test when significant differences were identified among the variances. A significant level of p<0.05 was established for the comparisons between groups. The statistical analyses were performed using the Prism Version 8.0.2 software package (Graph Pad Inc., USA).

## 3. Results

### 3.1 UHPLC-MS Chemical Profiling of *K. grandifoliola* Hydro-Ethanolic Extract (KHE)

The UHPLC-MS analysis of KHE revealed a diverse chemical composition, as depicted in the chromatogram (Fig. 1). Through alignment of retention times (Rt) and mass-to-charge (m/z) ratios with established databases, 18 distinct compounds were characterized, representing multiple phytochemical classes. The identified constituents comprised eight flavonoid-O-glycosides, three polyflavonoids, two quinic acid derivatives, along with single representatives of phenolic glycoside, coumarin glycoside, terpene glycoside, benzoic acid derivative, and fatty acid (Table 1). Notable peaks corresponded to Chlorogenic acid (Rt = 5.63 min), Scopolin (Rt = 6.48 min), Quercitrin (Rt = 7.81 min), Trimethyl 3-methoxybenzene-1,2,4-tricarboxylate (Rt = 8.09 min), Genistin (Rt = 8.45 min), and Apigenin 7-O-(6’’-O-acetylglucoside) (Rt = 9.21 min), which were among the most abundant. Additional significant compounds included kaempferol 3-galactoside-7-rhamnoside (Rt = 7.19 min), 3-O-p-coumaroylquinic acid (Rt = 7.25 min), and caffeic acid 3-O-glucuronide (Rt = 11.26 min), underscoring KHE’s complex phytochemical profile.

**Fig. 1:**
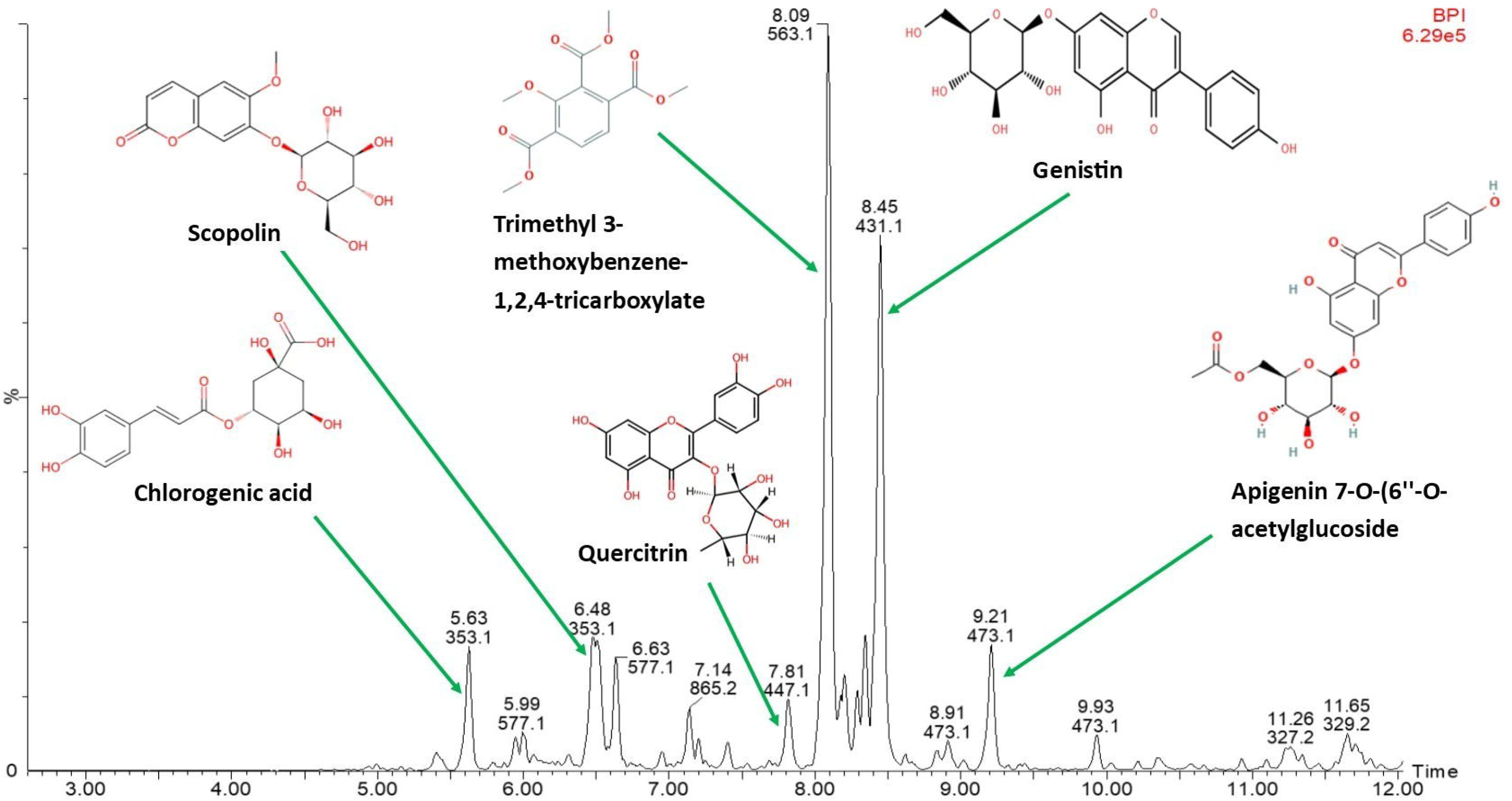
UHPLC-MS chromatographic profile of the hydro-ethanolic extract of *K. grandifoliola* (KHE) Major compounds identified: Chlorogenic acid (Rt = 5.63 min); Scopolin (Rt = 6.48 min); Quercitrin (Rt = 7.81 min); Trimethyl 3-methoxybenzene-1,2,4-tricarboxylate (Rt = 8.09 min); Genistin (Rt = 8.45 min) and Apigenin 7-O-(6’’-O-acetylglucoside) (Rt = 9.47 min).

**Fig. 2:**
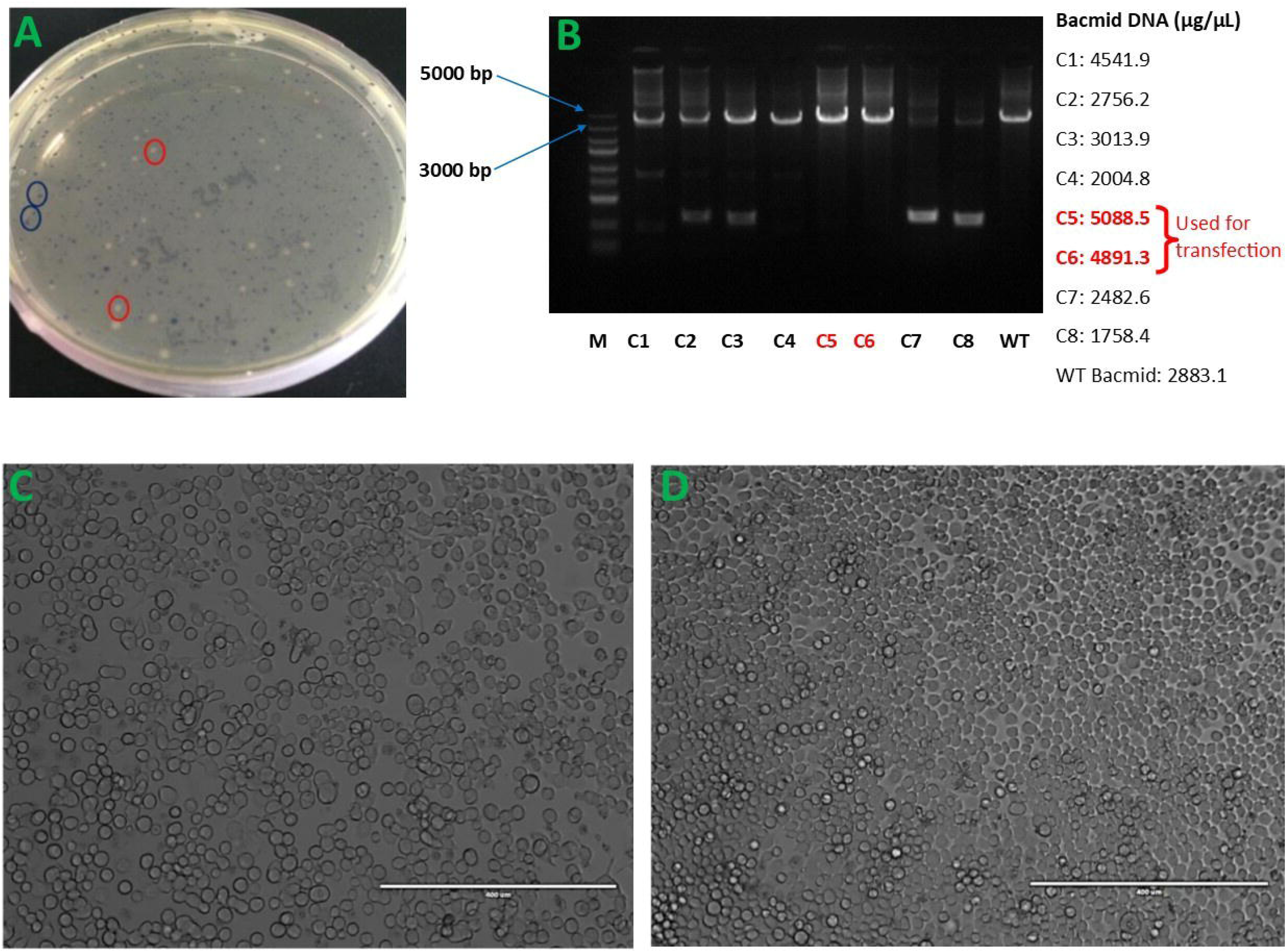
Recombinant DNA Bacmid and recombinant baculovirus carrying HEV-PLpro sequence (A): Selection of positive recombinant *DH*_α_*10Bac* colonies (white) harboring the HEV-PLpro expression construct (*pFastBac1-N-GP67-HEV-PLpro-C-6His*), alongside negative colonies (blue). **(B):** PCR verification of the correct insertion of the HEV-PLpro sequence (486 bp) into the Bacmid vector (4775 bp). **(C):** Morphology of *Sf9* cells at 72 h post-transfection with the recombinant Bacmid (10× magnification). **(D):** Morphology of non-transfected *Sf9* cells at 72 h (10× magnification).

**Table 1:**
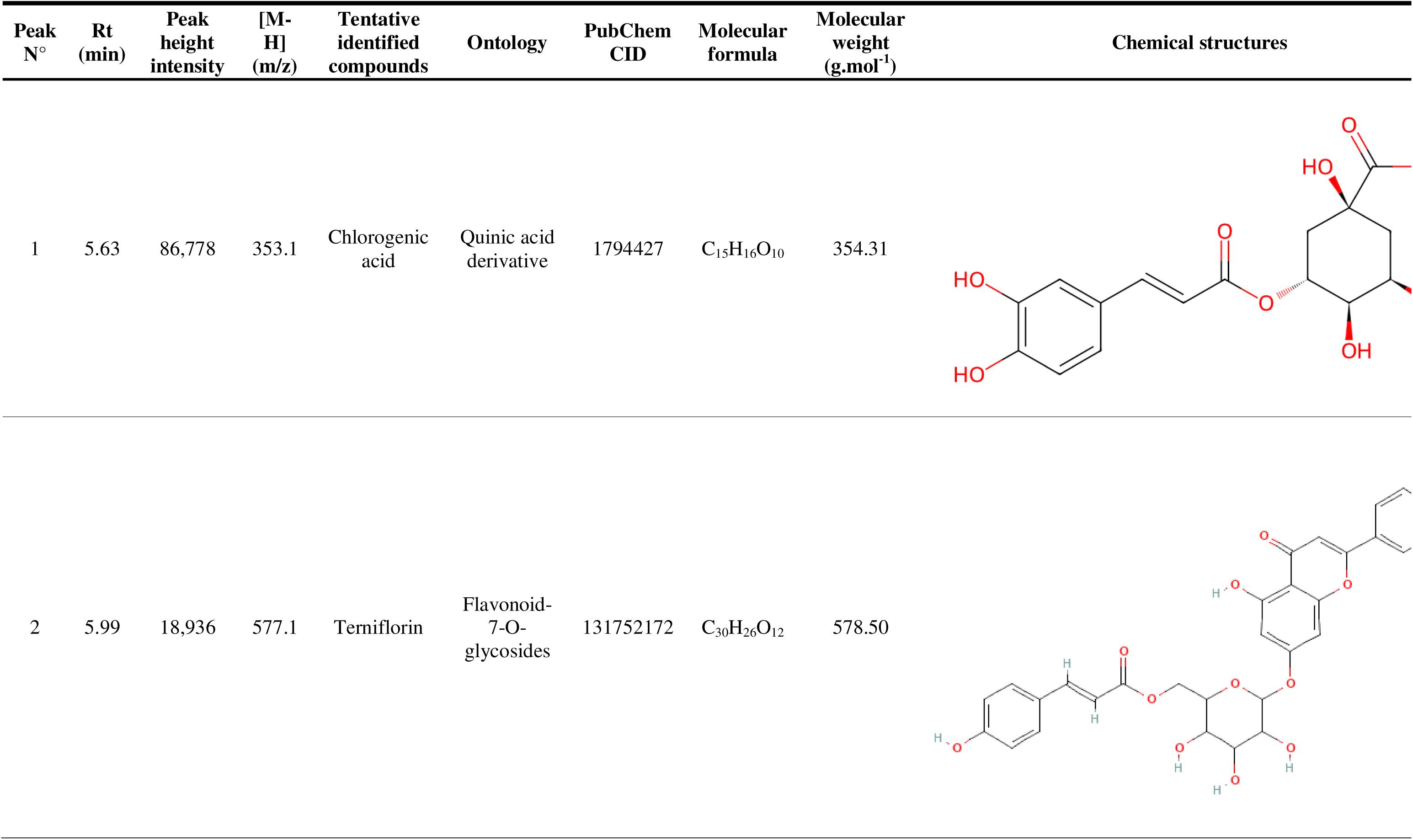
UHPLC-MS chemical profiling of the hydro-ethanolic extract of *K. grandifoliola* (Begin)

| Peak N° | Rt (min) | Peak height intensity | [M-H] (m/z) | Tentative identified compounds | Ontology | PubChem CID | Molecular formula | Molecular weight (g.mol<sup>-1</sup>) | Chemical structures |
| --- | --- | --- | --- | --- | --- | --- | --- | --- | --- |
| 1 | 5.63 | 86,778 | 353.1 | Chlorogenic acid | Quinic acid derivative | 1794427 | C<sub>15</sub>H<sub>16</sub>O<sub>10</sub> | 354.31 |  |
| 2 | 5.99 | 18,936 | 577.1 | Terniflorin | Flavonoid-7-O-glycosides | 131752172 | C<sub>30</sub>H<sub>26</sub>O<sub>12</sub> | 578.50 |  |

### 3.2 Generation of recombinant baculovirus harboring HEV-PLpro Sequence

Transformation of DHα10Bac competent cells with the HEV-PLpro expression construct (pFastBac1-N-GP67-HEV-PLpro-C-6His) yielded white recombinant colonies, distinguishable from non-recombinant blue clones (Fig. 2A). PCR-based screening further verified the correct integration of the HEV-PLpro insert (486 bp) into the Bacmid vector (4775 bp), as evidenced by a single ∼5100 bp amplicon in agarose gel electrophoresis (Fig. 2B). Subsequently, Sf9 cells were transfected with the recombinant Bacmid to generate the HEV-PLpro-expressing baculovirus. Successful transfection was confirmed by characteristic cytopathic effects, including vacuolization, granulation, and cellular enlargement (Fig. 2C), absent in non-transfected control cell (Fig. 2D).

### 3.3 Expression, purification, and enzymatic characterization of recombinant HEV-PLpro

High Five cells infected with a high-titer P4 baculovirus stock (amplified in Sf9 cells) expressed recombinant HEV-PLpro after 60 h. A two-step purification strategy starting with nickel-affinity chromatography (Fig. 3A) followed by size-exclusion chromatography yielded highly pure (>97%) HEV-PLpro, as confirmed by SDS-PAGE (Fig. 3B). The observed molecular weight (∼20 kDa) aligned closely with the predicted size (19.21 kDa). Enzymatic activity of the purified HEV-PLpro was subsequently assessed using the fluorogenic substrate Z-RLRGG-AMC. Michaelis-Menten kinetics (Fig. 3C) yielded approximate Vmax of 2.42 ± 0.1 µmol·minL¹ (hydrolyzed AMC) and Km of 28.79 ± 2.7 µM, while Lineweaver-Burk analysis (Fig. 3D) provided refined values of Vmax = 2.91 ± 0.3 µmol·minL¹ and Km = 32.05 ± 1.9 µM, demonstrating efficient proteolytic cleavage.

**Fig. 3:**
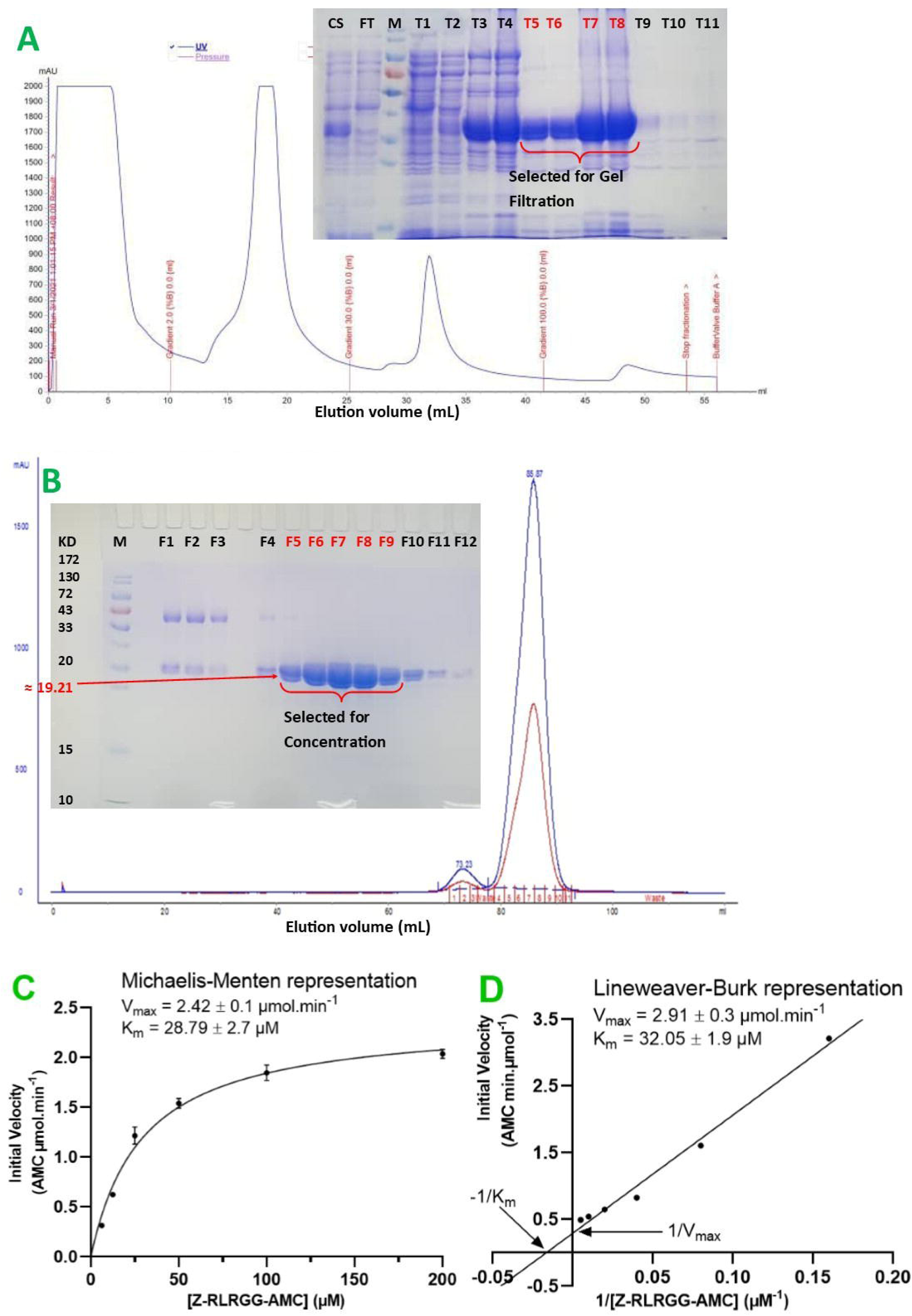
Expression, purification and enzyme kinetic parameters of the purified recombinant HEV-PLpro. HEV-PLpro was expressed in High Five cells through Baculovirus expression system and purified by affinity and size exclusion chromatography. Then, its proteolytic activity was evaluated using Z-RLRGG-AMC substrate. **(A)**: Chromatogram of affinity chromatography and SDS-PAGE analysis of the fractions obtained. **(B)**: Chromatogram of size exclusion chromatography and SDS-PAGE analysis of the fractions obtained. **(C)** and **(D)**: Michaelis-Menten and Lineweaver-Burk Plots for the proteolytic activity of the purified recombinant HEV-PLpro, respectively.

### 3.4 Concentration-dependent inhibition of HEV-PLpro proteolytic activity by KHE

The inhibitory effects of KHE and ribavirin (RBV), the reference inhibitor, on the proteolytic activity of HEV-PLpro were assessed across a range of concentrations. As illustrated in Fig. 4A, both tested inhibitors exhibited a concentration-dependent suppression of HEV-PLpro activity. The half-maximal inhibitory concentrations (IC_50_) for KHE and RBV were determined to be 20.86 ± 1.2 μg/mL and 19.13 ± 0.5 μg/mL, respectively. Although KHE displayed a marginally higher IC_50_ than RBV, statistical analysis revealed no significant difference between the two (P > 0.05), suggesting that KHE possesses inhibitory efficacy comparable to that of RBV.

**Fig. 4:**
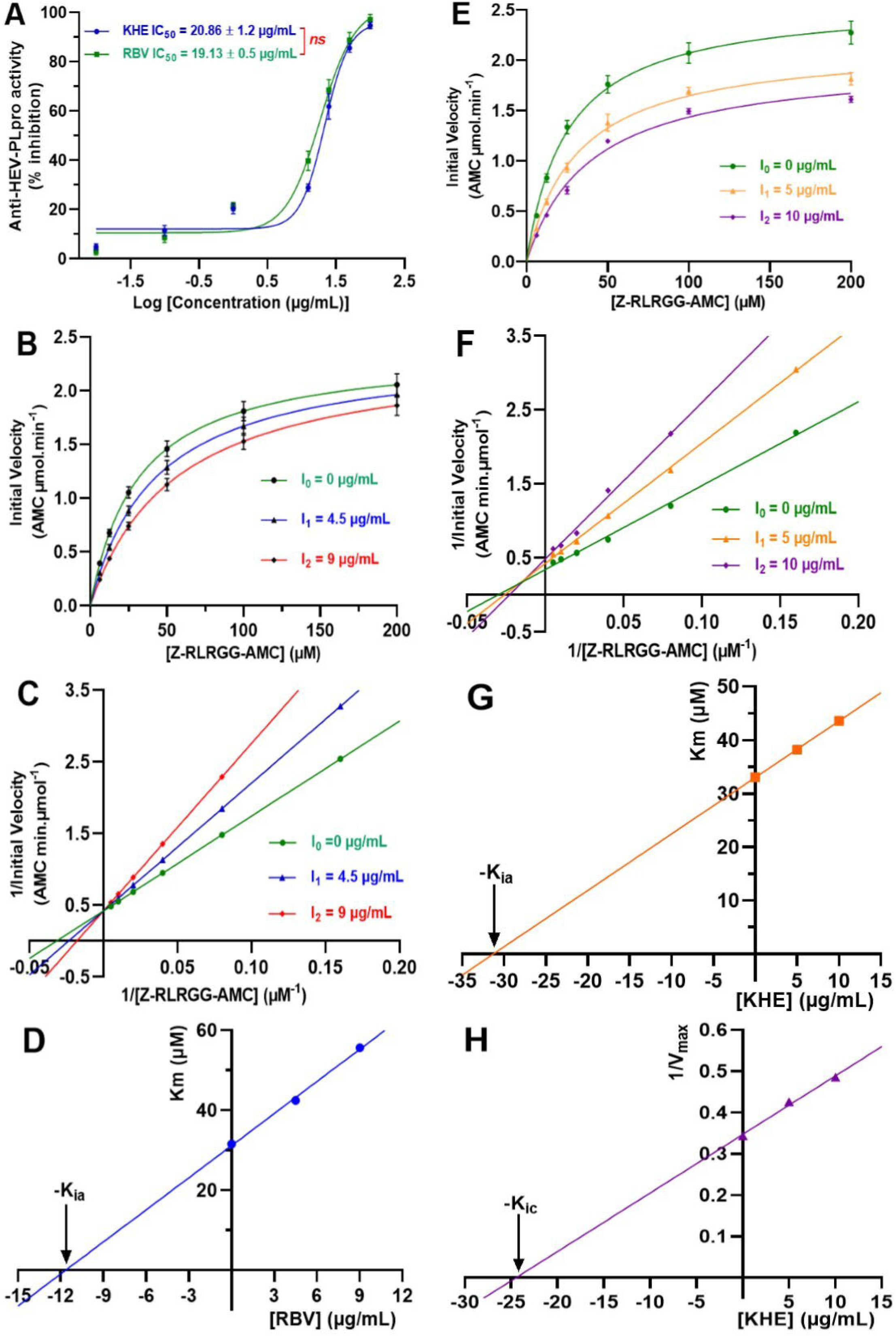
Inhibitory mechanism of KHE and RBV on the proteolytic activity of HEV-PLpro (A): Concentration-response curve showing the inhibitory effect of KHE and RBV on the proteolytic activity of HEV-PLpro. **(B)** and **(E)**: Michaelis-Menten plot of proteolytic activity of HEV-PLpro in presence of RBV and KHE respectively; **(C)** and **(F)**: Double-reciprocal plot of of proteolytic activity of HEV-PLpro in presence of RBV and KHE respectively; **(D)** and **(G)**: Secondary plot (K_m_’= f[I]) for the determination of the inhibition constant (K_ia_) for RBV and KHE respectively; **(H)**: Secondary plot (1/V_max_’= f[I]) for the determination of the inhibition constant (K_ic_) for KHE. KHE: Data are expressed as Means ± SD of three independent experiments in triplicate. ^ns^ Values non-significantly different (P>0.05) when compared using unpaired Student *t* test. KHE: Hydro-ethanolic extract of *K. grandifoliola*; RBV: Ribavirin; K_ia_: Inhibition constant toward K_m_; K_ic_: Inhibition constant toward V_max_.

### 3.5 Inhibition Mechanism of KHE on the proteolytic activity of HEV-PLpro

After determining the IC_50_ values, the inhibitory mechanisms of KHE and RBV were investigated. The initial reaction rates were measured at varying substrate concentrations, both in the presence and absence of inhibitors. Kinetic analysis included Michaelis-Menten plots (Figs. 4B and 4E for RBV and KHE, respectively), Lineweaver-Burk double reciprocal plots (Figs. 4C and 4F), and secondary plots of Km’ vs. [I] (Figs. 4D and 4G) and 1/Vmax’ vs. [I] (Fig. 4H). The derived kinetic parameters are summarized in Table 2.

**Table 2:**
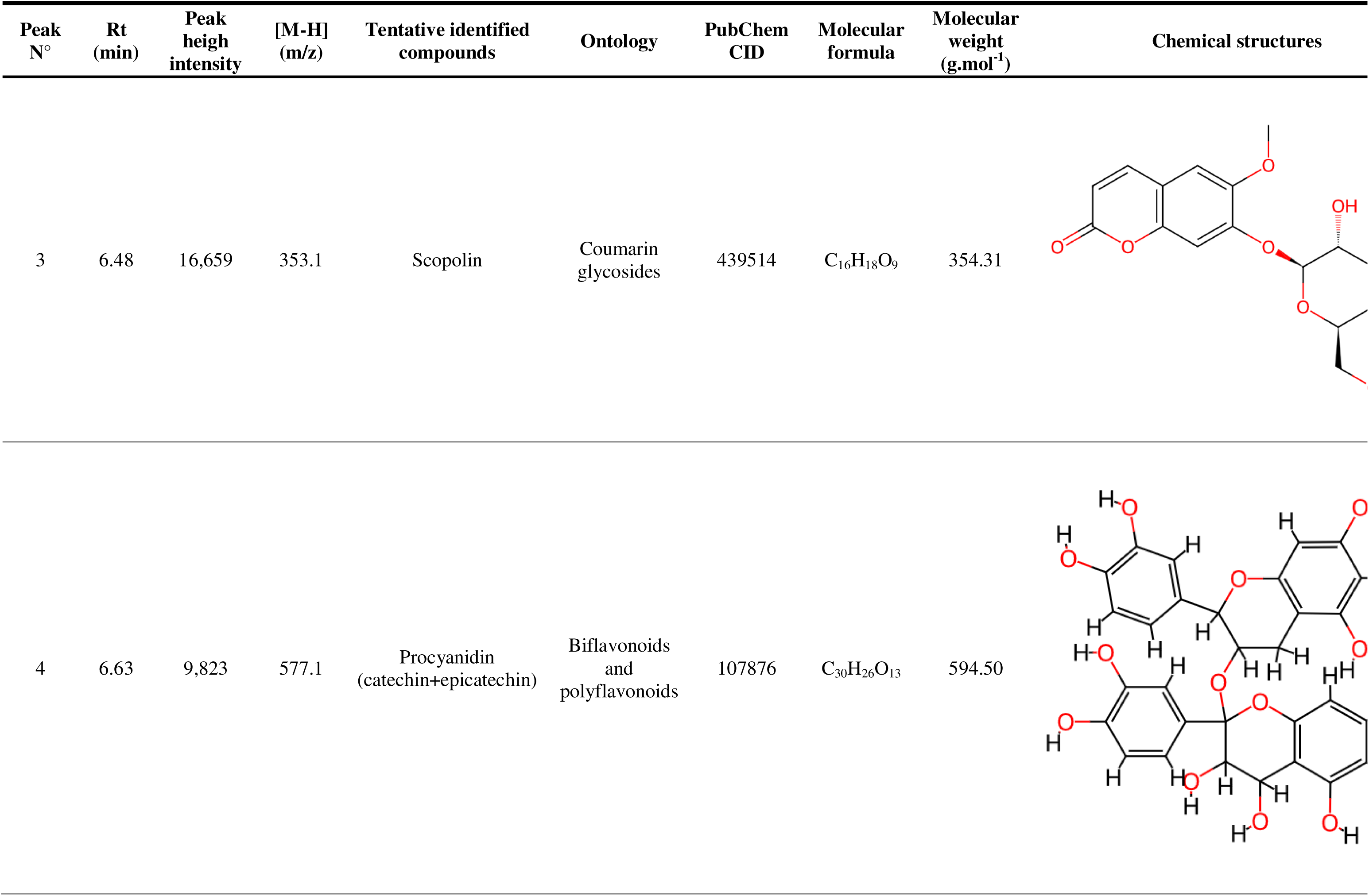
UHPLC-MS chemical profiling of the hydro-ethanolic extract of *K. grandifoliola* (Continued)

| Peak N° | Rt (min) | Peak heigh intensity | [M-H] (m/z) | Tentative identified compounds | Ontology | PubChem CID | Molecular formula | Molecular weight (g.mol<sup>-1</sup>) | Chemical structures |
| --- | --- | --- | --- | --- | --- | --- | --- | --- | --- |
| 3 | 6.48 | 16,659 | 353.1 | Scopolin | Coumarin glycosides | 439514 | C<sub>16</sub>H<sub>18</sub>O<sub>9</sub> | 354.31 |  |
| 4 | 6.63 | 9,823 | 577.1 | Procyanidin (catechin+epicatechin) | Biflavonoids and polyflavonoids | 107876 | C<sub>30</sub>H<sub>26</sub>O<sub>13</sub> | 594.50 |  |

RBV exhibited competitive inhibition, as evidenced by the Lineweaver-Burk plot, where all lines intersected at the same Y-intercept (Fig. 4C). The inhibition constant (*Kia*), reflecting RBV’s affinity for the free enzyme, was calculated as 11.63 ± 1.40 μg/mL from the secondary plot of Km’ vs. [I] (Fig. 4D). In contrast, KHE displayed mixed-type inhibition, indicated by converging lines in the Lineweaver-Burk plot that intersected to the left of the Y-axis and above the X-axis (Fig. 4F). Therefore, two distinct inhibition constants were derived: Kia = 31.28 ± 0.91 µg/mL (affinity for the free enzyme) obtained from Km’ vs. [I] (Fig. 4G); and Kic = 24.44 ± 1.61 µg/mL (affinity for the enzyme-substrate complex), obtained from 1/Vmax’ vs. [I] (Fig. 4H).

### 3.6 Molecular binding interactions between HEV-ORF1 and KHE-derived compounds

To further elucidate the potential molecular mechanism underlying the inhibitory effect of KHE against HEV-PLpro activity, molecular docking was conducted between the HEV-ORF1 protein (PDB ID: 6NU9), which harbors the papain-like protease (PLpro) domain, and the phytochemicals identified in KHE via UHPLC-MS analysis. The protein-ligand binding affinities were evaluated based on docking scores (kcal/mol). Among the 18 screened compounds, six exhibited superior binding affinities compared to the reference inhibitor, ribavirin (RBV). In particular, scopolin (L3), procyanidin (L3), quercetin 3-[rhamnosyl-(1→2)-α-L-arabinopyranoside] (L10), and quercitrin (L12) with their 3D and 2D interaction profiles with HEV-ORF1 illustrated by Fig. 5, demonstrated the strongest interactions, with docking scores of −7.07, −6.23, −7.68, and −7.43 kcal/mol, respectively, significantly outperforming RBV (−4.87 kcal/mol) (Table 3). The most promising ligand, L10, formed ten hydrogen bonds with key residues (His164, Glu166, Glu182, Pro177, Asp55, Ser68, and Gly178), along with three hydrophobic interactions (two Pi-alkyl bonds with Ala73 and Pro177, and one Pi-sigma bond with Arg66). In contrast, RBV established five conventional hydrogen bonds with Asp55, Arg66, Thr67, and Trp181, three carbon-hydrogen bonds with Ser68, His179, and Trp181), one amide-Pi stacking and one Pi-alkyl interaction with Thr67 and Arg66, respectively (Table 3).

**Fig. 5:**
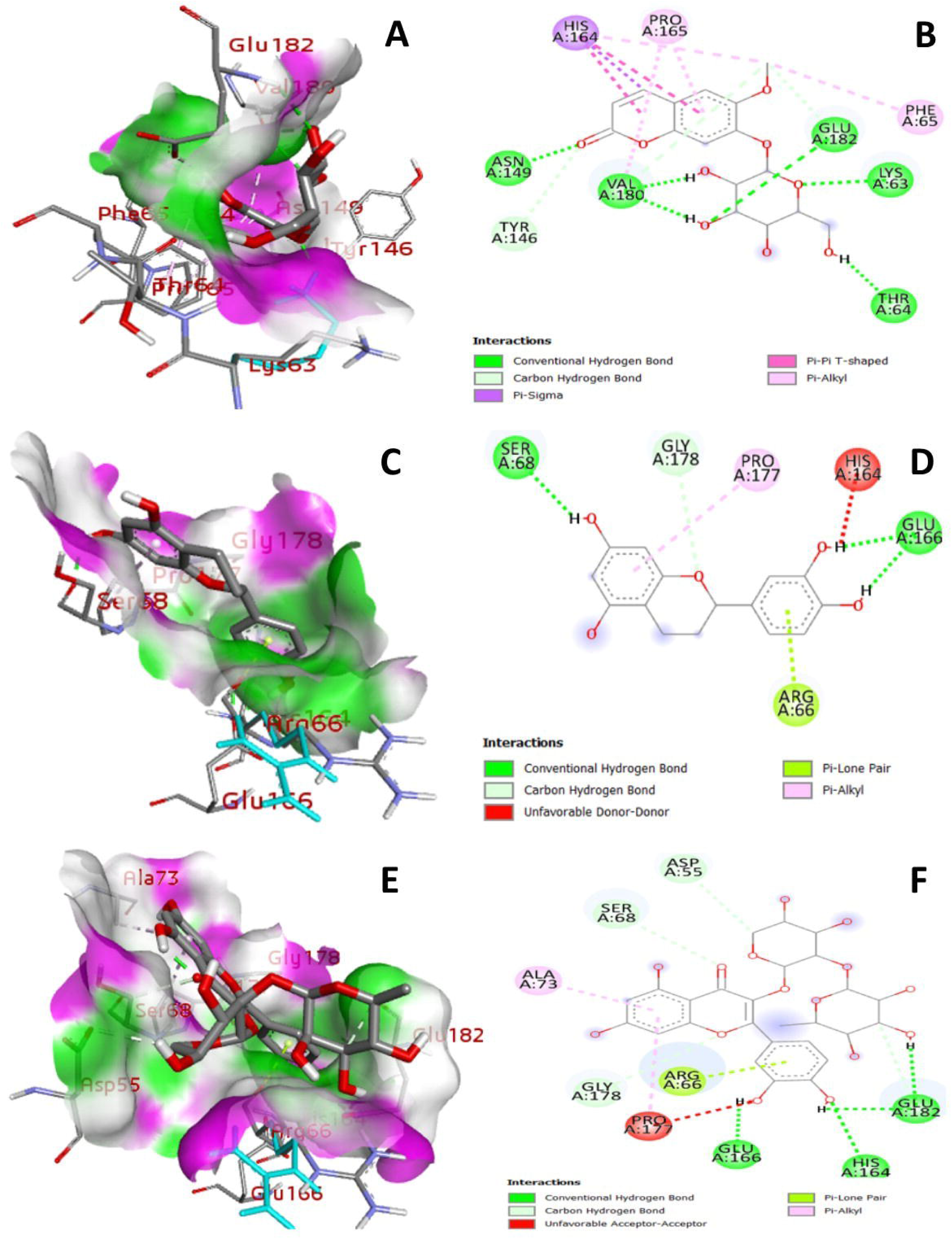
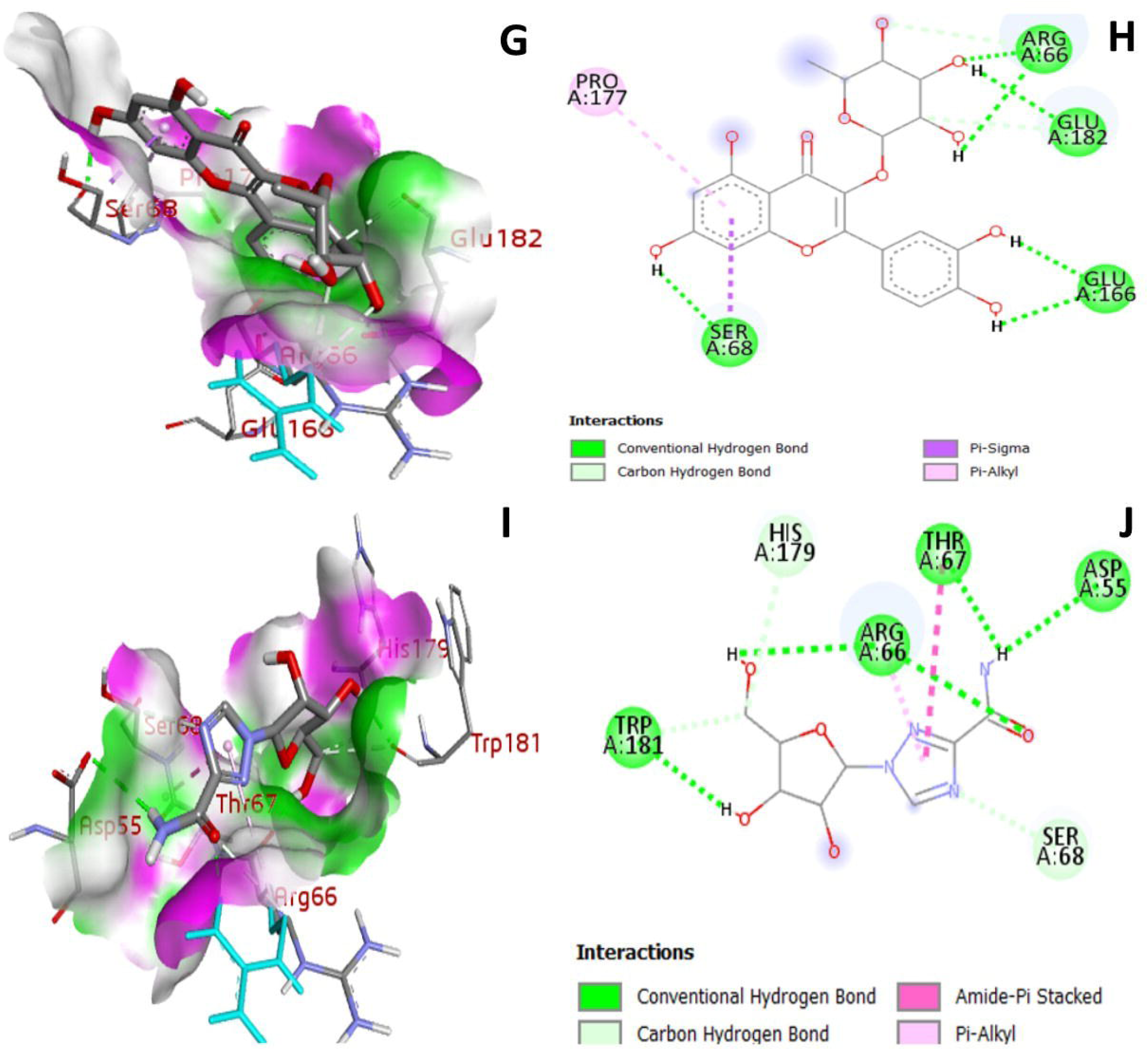
Visualized 3-D and 2-D protein-ligand complexes between HEV_ORF1 (PDB ID: 6NU9) and the hit compounds identified from KHE. **(A)**, **(C)**, **(E)**, **(G)**, and **(I)**: 3D receptor-ligand view between HEV_ORF1 and Scopolin, Procyanidin, Quercetin 3-[rhamnosyl-(1->2)-alpha-L-arabinopyranoside], Quercitrin, and Ribavirin, respectively. **(B)**, **(D)**, **(F)**, **(H)**, and **(J)**: 2D receptor-ligand view showing molecular interactions between HEV_ORF1 and Scopolin, Procyanidin, Quercetin 3-[rhamnosyl-(1->2)-alpha-L-arabinopyranoside], Quercitrin, and Ribavirin, respectively.

**Table 3:**
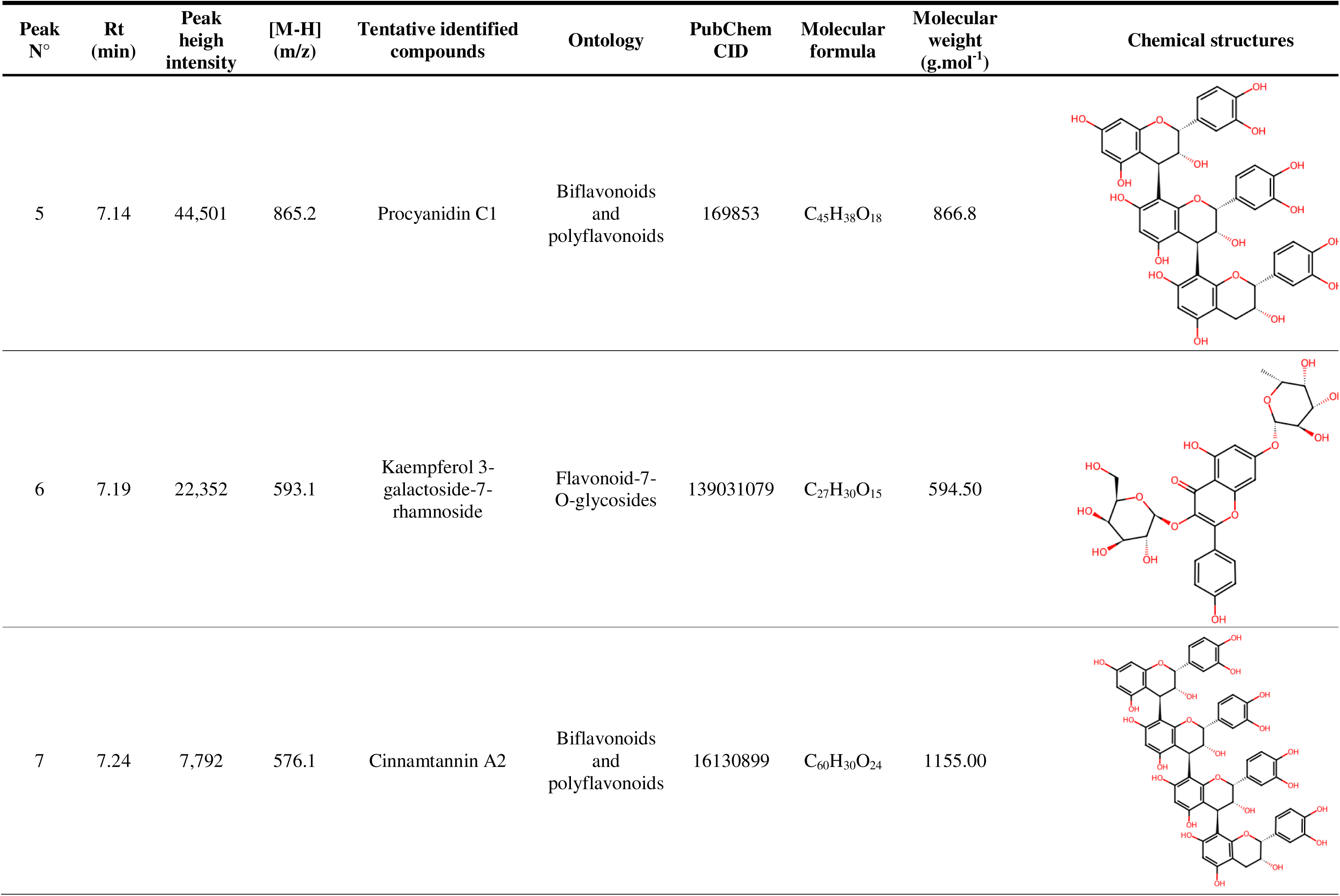
UHPLC-MS chemical profiling of the hydro-ethanolic extract of *K. grandifoliola* (Continued)

| Peak N° | Rt (min) | Peak heigh intensity | [M-H] (m/z) | Tentative identified compounds | Ontology | PubChem CID | Molecular formula | Molecular weight (g.mol<sup>-1</sup>) | Chemical structures |
| --- | --- | --- | --- | --- | --- | --- | --- | --- | --- |
| 5 | 7.14 | 44,501 | 865.2 | Procyanidin C1 | Biflavonoids and polyflavonoids | 169853 | C<sub>45</sub>H<sub>38</sub>O<sub>18</sub> | 866.8 |  |
| 6 | 7.19 | 22,352 | 593.1 | Kaempferol 3-galactoside-7-rhamnoside | Flavonoid-7-O-glycosides | 139031079 | C<sub>27</sub>H<sub>30</sub>O<sub>15</sub> | 594.50 |  |
| 7 | 7.24 | 7,792 | 576.1 | Cinnamtannin A2 | Biflavonoids and polyflavonoids | 16130899 | C<sub>60</sub>H<sub>30</sub>O<sub>24</sub> | 1155.00 |  |

**Table 4:**
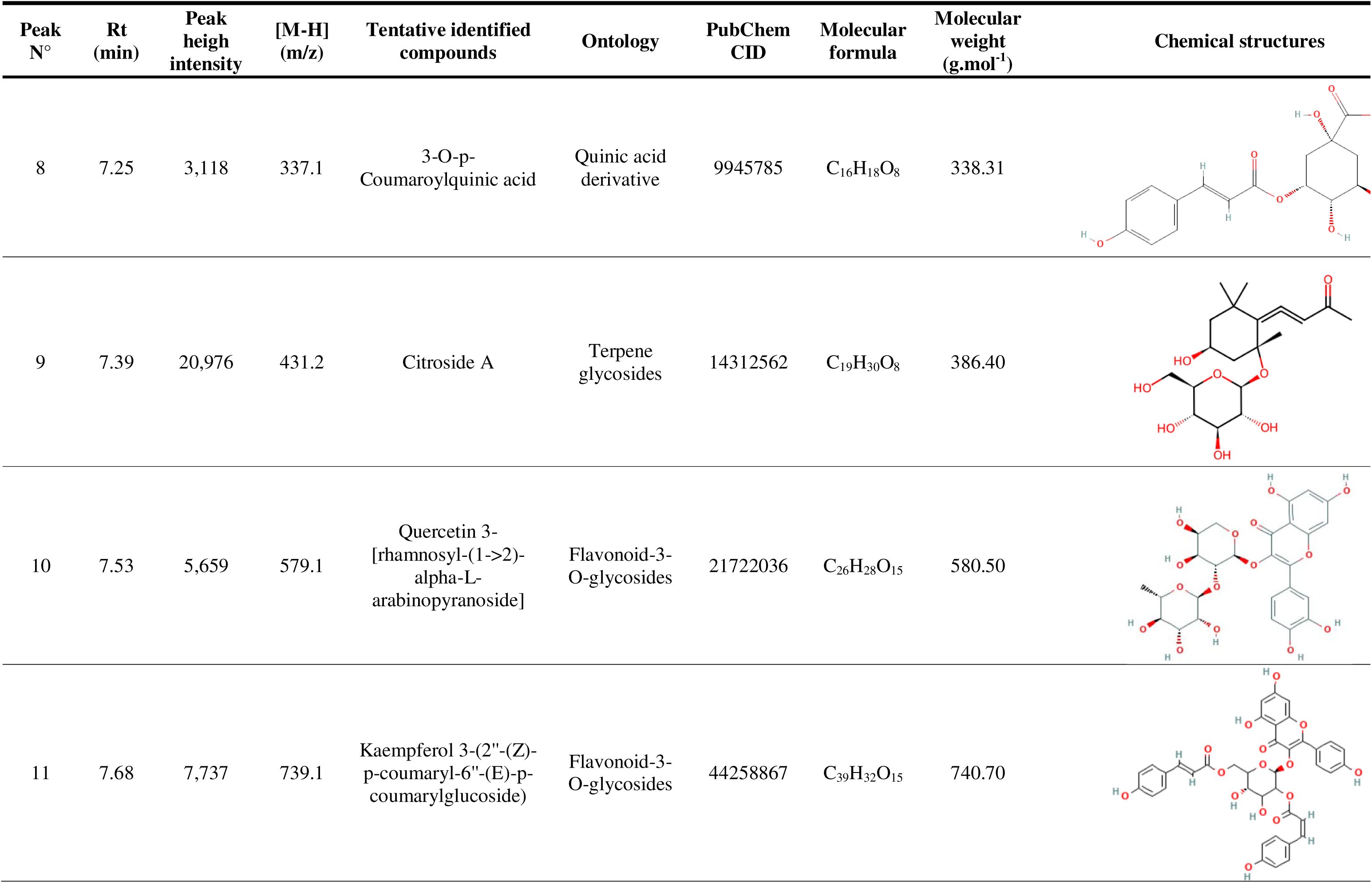
UHPLC-MS chemical profiling of the hydro-ethanolic extract of *K. grandifoliola* (Continued)

| Peak N° | Rt (min) | Peak heigh intensity | [M-H] (m/z) | Tentative identified compounds | Ontology | PubChem CID | Molecular formula | Molecular weight (g.mol<sup>-1</sup>) | Chemical structures |
| --- | --- | --- | --- | --- | --- | --- | --- | --- | --- |
| 8 | 7.25 | 3,118 | 337.1 | 3-O-p-Coumaroylquinic acid | Quinic acid derivative | 9945785 | C<sub>16</sub>H<sub>18</sub>O<sub>8</sub> | 338.31 |  |
| 9 | 7.39 | 20,976 | 431.2 | Citroside A | Terpene glycosides | 14312562 | C<sub>19</sub>H<sub>30</sub>O<sub>8</sub> | 386.40 |  |
| 10 | 7.53 | 5,659 | 579.1 | Quercetin 3-[rhamnosyl-(1->2)-alpha-L-arabinopyranoside] | Flavonoid-3-O-glycosides | 21722036 | C<sub>26</sub>H<sub>28</sub>O<sub>15</sub> | 580.50 |  |
| 11 | 7.68 | 7,737 | 739.1 | Kaempferol 3-(2''-(Z)-p-coumaryl-6''-(E)-p-coumarylglucoside) | Flavonoid-3-O-glycosides | 44258867 | C<sub>39</sub>H<sub>32</sub>O<sub>15</sub> | 740.70 |  |

**Table 5:**
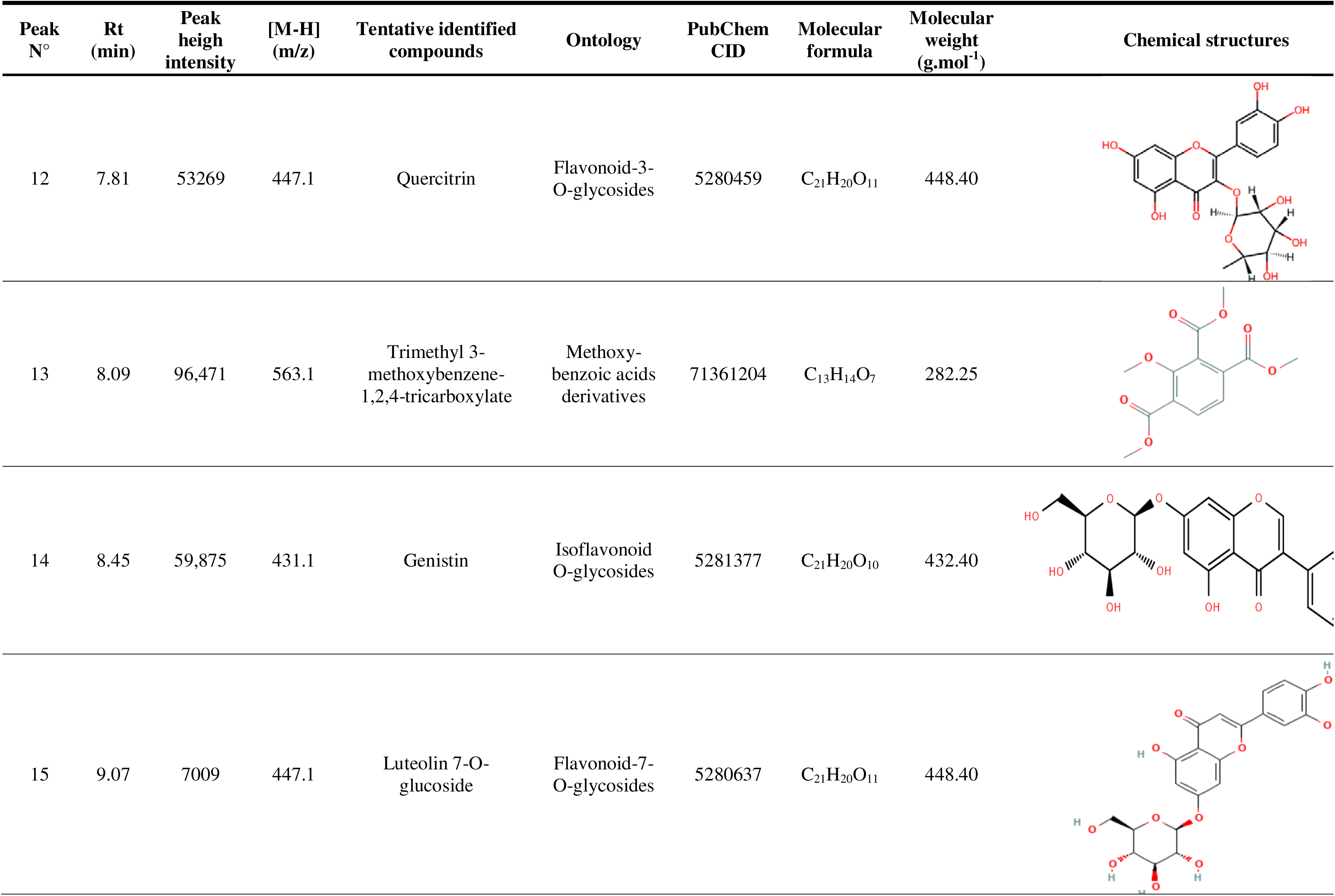
UHPLC-MS chemical profiling of the hydro-ethanolic extract of *K. grandifoliola* (Continued)

| Peak N° | Rt (min) | Peak heigh intensity | [M-H] (m/z) | Tentative identified compounds | Ontology | PubChem CID | Molecular formula | Molecular weight (g.mol<sup>-1</sup>) | Chemical structures |
| --- | --- | --- | --- | --- | --- | --- | --- | --- | --- |
| 12 | 7.81 | 53269 | 447.1 | Quercitrin | Flavonoid-3-O-glycosides | 5280459 | C<sub>21</sub>H<sub>20</sub>O<sub>11</sub> | 448.40 |  |
| 13 | 8.09 | 96,471 | 563.1 | Trimethyl 3-methoxybenzene-1,2,4-tricarboxylate | Methoxy-benzoic acids derivatives | 71361204 | C<sub>13</sub>H<sub>14</sub>O<sub>7</sub> | 282.25 |  |
| 14 | 8.45 | 59,875 | 431.1 | Genistin | Isoflavonoid O-glycosides | 5281377 | C<sub>21</sub>H<sub>20</sub>O<sub>10</sub> | 432.40 |  |
| 15 | 9.07 | 7009 | 447.1 | Luteolin 7-O-glucoside | Flavonoid-7-O-glycosides | 5280637 | C<sub>21</sub>H<sub>20</sub>O<sub>11</sub> | 448.40 |  |

**Table 6:**
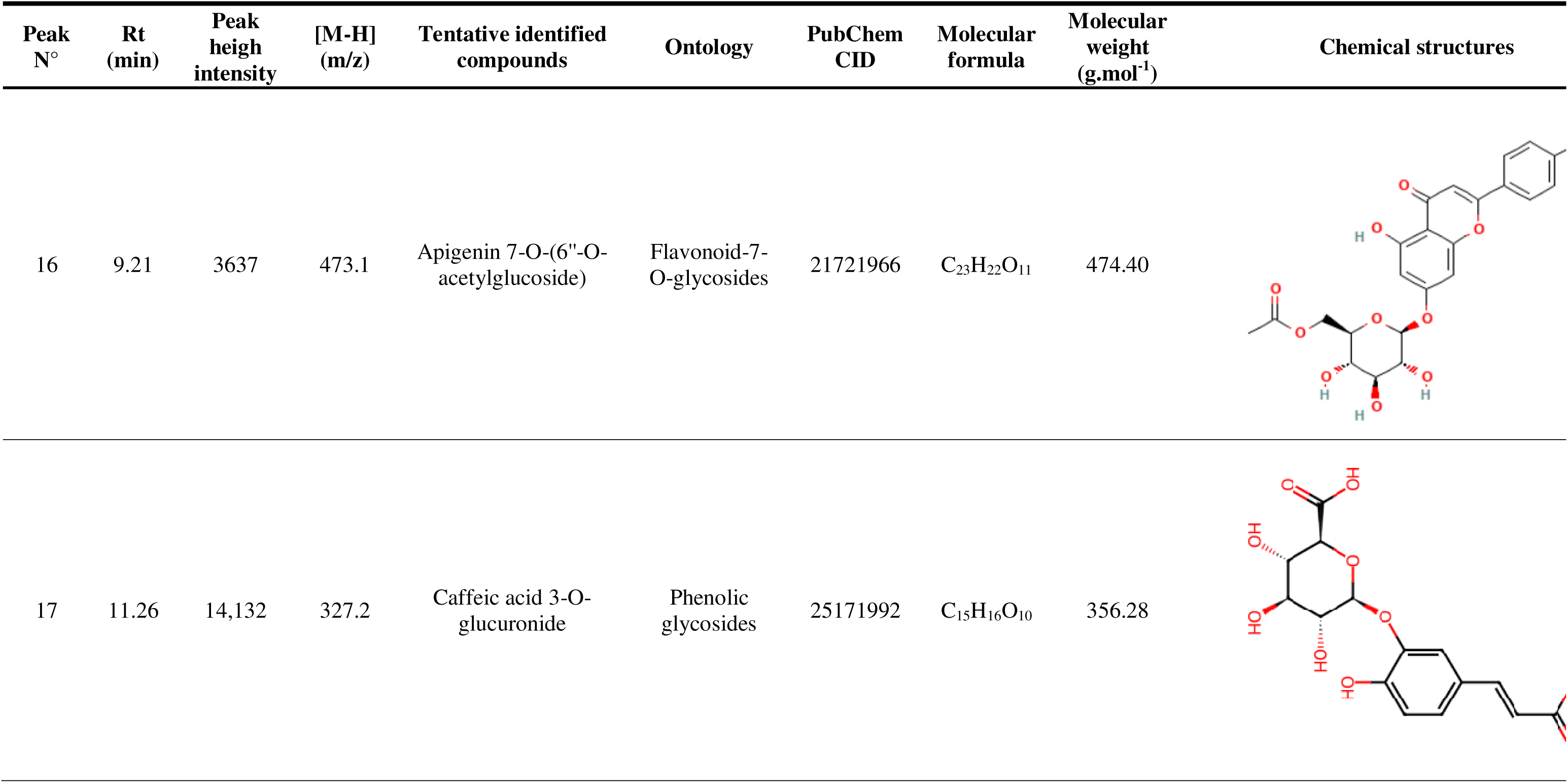
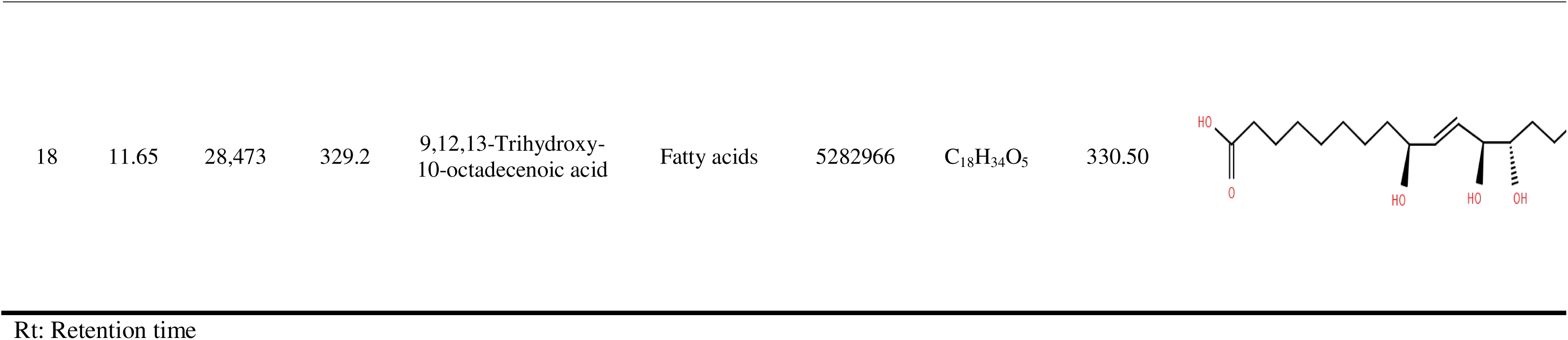
UHPLC-MS chemical profiling of the hydro-ethanolic extract of *K. grandifoliola* (End)

| Peak N° | Rt (min) | Peak heigh intensity | [M-H] (m/z) | Tentative identified compounds | Ontology | PubChem CID | Molecular formula | Molecular weight (g.mol<sup>-1</sup>) | Chemical structures |
| --- | --- | --- | --- | --- | --- | --- | --- | --- | --- |
| 16 | 9.21 | 3637 | 473.1 | Apigenin 7-O-(6"-O-acetylglucoside) | Flavonoid-7-O-glycosides | 21721966 | C<sub>23</sub>H<sub>22</sub>O<sub>11</sub> | 474.40 |  |
| 17 | 11.26 | 14,132 | 327.2 | Caffeic acid 3-O-glucuronide | Phenolic glycosides | 25171992 | C<sub>15</sub>H<sub>16</sub>O<sub>10</sub> | 356.28 |  |

|  |  |  |  |  |  |  |  |  |
| --- | --- | --- | --- | --- | --- | --- | --- | --- |
| 18 | 11.65 | 28,473 | 329.2 | 9,12,13-Trihydroxy-10-octadecenoic acid | Fatty acids | 5282966 | C<sub>18</sub>H<sub>34</sub>O<sub>5</sub> | 330.50 |

Rt: Retention time

### 3.7 Molecular Dynamics Simulation Analysis

To further validate the stability and structural dynamics of the top-performing HEV-ORF1 (PDB ID: 6NU9) complexes identified through docking, molecular dynamics simulations (MDS) were conducted over 100 ns for the L10-6NU9 and L12-6NU9 complexes, alongside the reference inhibitor RBV-6NU9. Key parameters, including root-mean-square deviation (RMSD), root-mean-square fluctuation (RMSF), radius of gyration (Rg), and solvent-accessible surface area (SASA), were analyzed to assess conformational stability, flexibility, compactness, and solvent exposure.

#### 3.7.1 RMSD Analysis

The RMSD of the protein backbone atoms was calculated to evaluate structural deviations over time. The L12-6NU9 complex exhibited the lowest average RMSD (1.93 Å), followed by RBV-6NU9 (2.22 Å) and L10-6NU9 (2.33 Å) (Fig. 6A), indicating superior stability of the L12-bound system. Minimal fluctuations across all trajectories suggested sustained structural integrity throughout the simulation.

**Fig. 6:**
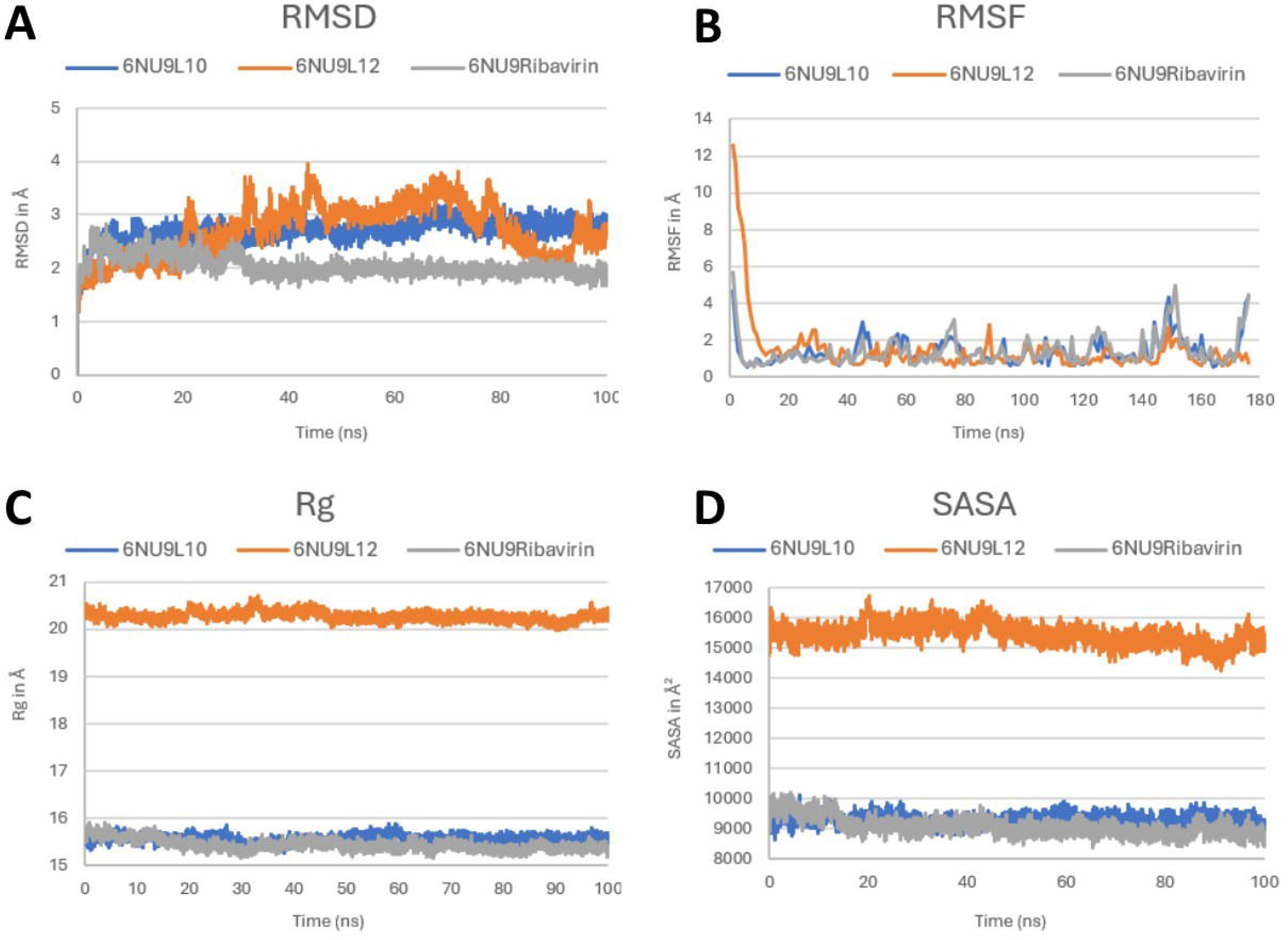
Molecular dynamics simulation data for the top two hits complexes and reference inhibitor. **(A)**: Root means square deviation (RMSD). **(B)**: Root means square fluctuation (RMSF). **(C)**: Radius of gyration (Rg). **(D)**: Solvent accessible surface area (SASA). 6NU9-L10: HEV_ORF1 in complex with Quercetin 3-[rhamnosyl-(1->2)-alpha-L-arabinopyranoside]. 6NU9-L12: HEV_ORF1 in complex with Quercitrin. 6NU9-ribavirin: HEV_ORF1 in complex with ribavirin.

#### 3.7.2 RMSF Analysis

Residue-specific flexibility was assessed via RMSF, revealing average fluctuations of 1.39 Å (L10), 1.47 Å (L12), and 1.41 Å (RBV) (Fig. 6B). Notably, all complexes displayed low RMSF values, signifying restricted residue mobility and stable ligand binding. No significant destabilizing peaks were observed, further supporting robust interactions.

#### 3.7.3 Radius of Gyration (Rg) Analysis

The Rg metric, reflecting tertiary structure compactness, yielded averages of 15.58 Å (L10), 20.28 Å (L12), and 15.65 Å (RBV) (Fig. 6C). While L12 induced a marginally higher Rg, the narrow deviation range across systems suggested no major structural unfolding, consistent with maintained protein stability.

#### 3.7.4 SASA Analysis

SASA measurements quantified solvent exposure, with averages of 9300.74 Å² (L10), 15450.9 Å² (L12), and 9085.58 Å² (RBV) (Fig. 6D). The elevated SASA for L12 implied slight surface exposure, yet the overall trends aligned with stable folding, as no abrupt changes occurred during simulation.

### 3.8 Anti-HEV Replication Activity and Cytotoxicity of KHE

The anti-HEV replication activity of KHE evaluated using a bio-luminescent reporter virus assay, revealed a concentration-dependent inhibition of viral replication. Notably, KHE (IC_50_ = 12.75 ± 1.9 µg/mL) demonstrated significantly stronger inhibitory potency (P < 0.05) than the reference drug ribavirin (RBV; IC_50_ = 17.96 ± 1.2 µg/mL) (Fig. 7A).

**Fig. 7:**
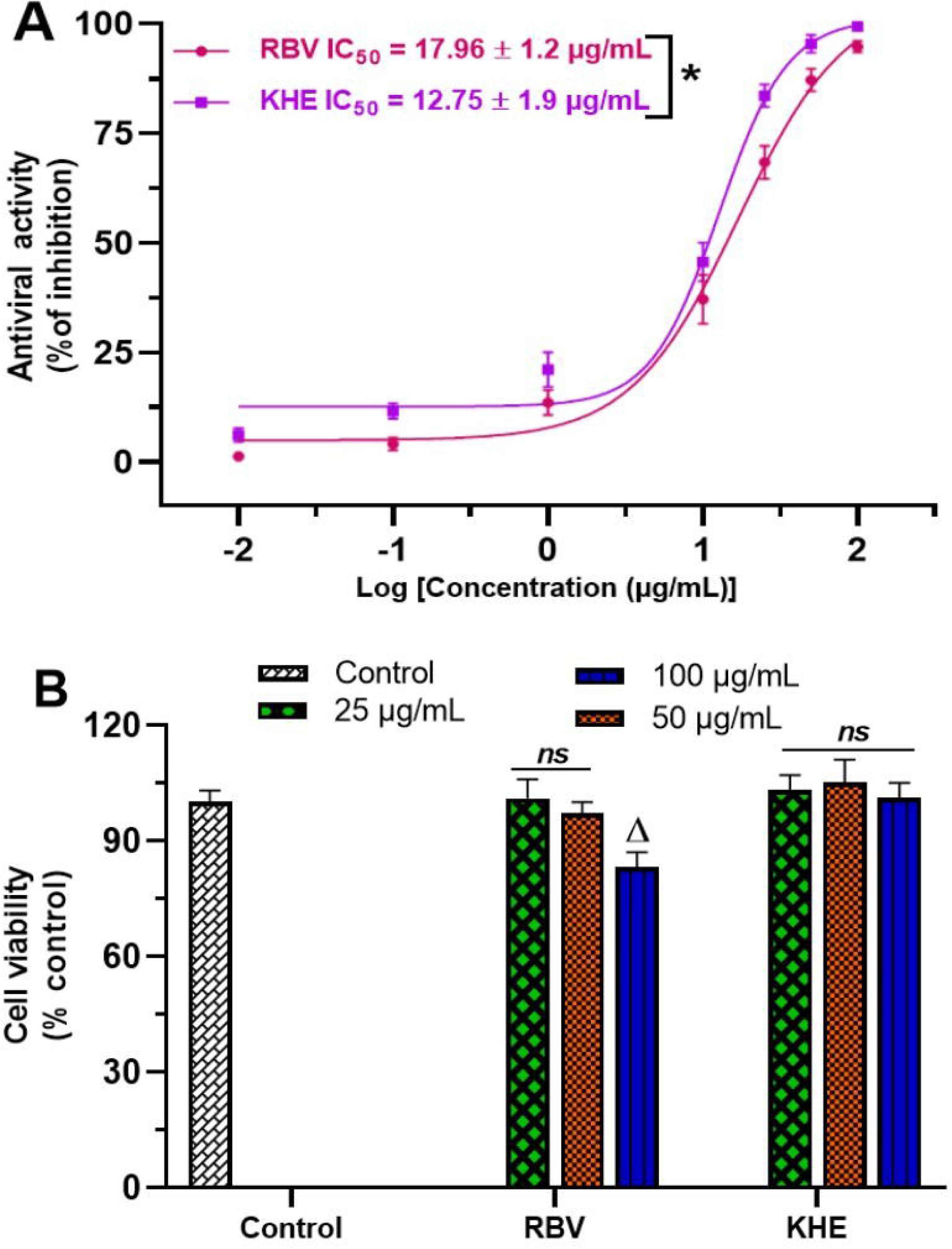
Anti-HEV replication activity and cytotoxicity effect of KHE (A): Concentration-response curve showing the inhibitory effect of KHE and RBV on the HEV replication. **(B)**: cytotoxicity effect of KHE and RBV on Hu7.5 cells. Data are expressed as Means ± SD of three independent experiments in triplicate. **\*** Values significantly different (P<0.05) when compared using unpaired Student *t* test. **^ns^** Values non-significantly different (P>0.05) when compared to the control cells using one-way ANOVA followed by Bonferoni’s post hoc test. ^Δ^ Values significantly different (P<0.05) when compared to the control cells using one-way ANOVA followed by Bonferoni’s post hoc test. KHE: Hydro-ethanolic extract of *K. grandifoliola*; RBV: Ribavirin.

Furthermore, cytotoxicity assays on Huh7.5 cells indicated that KHE exhibited no significant reduction in cell viability even at high concentrations (up to 100 µg/mL after 72 h). In contrast, RBV caused a marked decrease (P < 0.05) in cell viability at 100 µg/mL compared to untreated controls (Fig. 7B), highlighting KHE’s superior safety profile.

## 4. Discussion

Hepatitis E virus (HEV) infection remains a major global health concern, contributing to severe liver disease and thousands of annual fatalities (WHO, 2025). Despite its significant impact, HEV remains understudied, leaving a critical gap in targeted antiviral treatments. Currently, therapeutic options are limited to pegylated interferon-alpha (PEG-IFNα) combined with ribavirin (RBV), which shows only ∼70% efficacy (Gorris et al., 2021). Moreover, this regimen is often linked to adverse side effects and the emergence of drug-resistant viral strains (Davoodi et al., 2018; Debing et al., 2016; Sulkowski et al., 2011; Wu et al., 2016). Accordingly, there is a need to identify safe and effective antiviral agents capable of treating both acute and chronic HEV infections while preventing disease progression. To address this, the present study explores the inhibitory potential of *Khaya grandifoliola* hydro-ethanolic extract (KHE) against hepatitis E virus papain-like protease (HEV-PLpro), a key viral enzyme. Through an integrated approach, including phytochemical profiling, enzymatic kinetics, computational modeling, and *in vitro* antiviral assays, the findings provide comprehensive mechanistic insights into the inhibitory effects of KHE against HEV infection.

It is well recognized that viral proteases are pivotal for the replication and pathogenesis of numerous viruses, including coronaviruses and hepatitis viruses. For instance, papain-like and chymotrypsin-like proteases, as well as the HCV NS3/4A protease, are indispensable for the life cycle of coronaviruses and hepatitis C virus (HCV), respectively (Butt et al., 2022; Gao et al., 2021; Kumar et al., 2016; Lei et al., 2014). Similarly, the hepatitis E virus (HEV) papain-like cysteine protease (HEV-PLpro), derived from the non-structural polyprotein encoded by ORF1, plays a crucial role in viral maturation and immune evasion (Kenney and Meng, 2019; Wang and Meng, 2021). By processing viral polyproteins into functional units, HEV-PLpro ensures the production and maturation of infectious virions (Glitscher et al., 2024; LeDesma et al., 2019). Additionally, it subverts host immunity by suppressing type I interferon responses (Kim and Myoung, 2018; Kim et al., 2016), making it an attractive antiviral target against HEV infection.

HEV remains a poorly studied virus, one of the major obstacles hindering the development of antiviral treatments for HEV infection has been the challenge in purifying active forms of viral enzymes, necessary for early-stage drug screening (Netzler et al., 2019). This limitation was addressed in the present study, where recombinant HEV-PLpro, successfully expressed using a baculovirus expression system (Fig. 2) and purified via chromatography techniques (Fig. 3), was employed to test the inhibitory potential of KHE against HEV. The results revealed that KHE concentration-dependently suppresses HEV-PLpro’s proteolytic activity, exhibiting an IC_50_ similar to ribavirin (RBV), used as positive control (Fig. 4A). Through the inhibition of HEV-PLpro activity, these findings indicate that KHE could likely hinder HEV replication by preventing the cleavage of viral polyproteins into functional components, ultimately reducing the generation and maturation of new virions.

To elucidate the inhibitory mechanisms of KHE on HEV-PLpro’s proteolytic activity, enzyme kinetics assays were conducted using varying substrate concentrations in the presence or absence of inhibitors (KHE or RBV). The inhibition mode and constants were determined using Lineweaver-Burk plots and secondary representations, respectively. For RBV, the Lineweaver-Burk plot revealed a series of lines intersecting at the same point on the Y-axis (Fig. 4C), indicating that RBV does not alter the maximum reaction rate (V_max_) but increases the Michaelis-Menten constant (K_m_) value (Table 2). This suggests a competitive inhibition mechanism, with a K_ia_ value of 11.63 µg/mL (Fig. 4D; Table 2). RBV likely competes with the substrate for the active site or binds to an allosteric site, inducing conformational changes that reduce substrate affinity and impair HEV-PLpro activity. In contrast, KHE exhibited a mixed-type inhibition mechanism. The Lineweaver-Burk plot showed lines intersecting to the left of the Y-axis and above the X-axis (Fig. 4F), demonstrating that KHE affects both substrate affinity (K_m_) and enzyme-substrate complex formation (V_max_). The deduced inhibition constants, K_ia_ (31.28 µg/mL) and K_ic_ (24.44 µg/mL), further support this dual inhibitory action. Unlike RBV, whose competitive inhibition can be overcome by excess substrate, KHE’s competence to bind both the free enzyme and enzyme-substrate complex provides a potential therapeutic advantage.

Indeed, UHPLC-MS analysis identified multiple bioactive compounds in KHE, including flavonoids-O-glycosides, polyflavonoids, and phenolic and terpenoid derivatives, known for their antiviral and enzyme-inhibitory properties (Lin et al., 2014; Tabari et al., 2021). Peculiarly, Quercetin 3-[rhamnosyl-(1->2)-alpha-L-arabinopyranoside] (L10), Quercitrin (L12), and Apigenin 7-O-(6’’-O-acetylglucoside) (L16), previously reported to inhibit viral proteases in coronavirus and HCV (Bhattacharya et al., 2022; Lin et al., 2014; Mandal et al., 2021), were detected. Their presence in KHE may explain the mixed-type inhibition, where some compounds may interfere with substrate binding while others impair catalytic efficiency, collectively offering broader inhibitory action.

Molecular docking analysis reinforced these conclusions by showing high binding affinities of KHE’s compounds to HEV-ORF1 protein. Especially, Quercetin-3-[rhamnosyl-(1->2)-alpha-L-arabino-pyranoside] and Quercitrin demonstrated superior binding energies (−7.68 and −7.43 kcal/mol, respectively) compared to RBV (−4.87 kcal/mol) (Table 3), suggesting their enhanced potential to interact with key catalytic residues of HEV-PLpro. Indeed, detailed interaction analysis showed that the top ligand formed: four conventional hydrogen bonds with His164, Glu166, and Glu182, five carbon-hydrogen bonds with Asp55, Ser68, Gly178, and Glu182, three hydrophobic interactions (Pi-alkyl with Ala73/Pro177; Pi-sigma with Arg66), and one unfavorable donor-donor bond with Pro177. These interactions (Fig. 5; Table 3), known to stabilize protein-ligand complexes (Kouam et al., 2025a, 2025b, 2024a), suggest dual binding of KHE’s compounds at both the active and allosteric sites of HEV-PLpro, consistent with the observed mixed inhibition pattern.

Molecular dynamics simulations provided additional validation of the stable binding between HEV-ORF1 and both Quercetin 3-[rhamnosyl-(1→2)-α-L-arabinopyranoside] and Quercitrin. The low root-mean-square deviation (RMSD <2.5 Å) (Fig. 6A) and fluctuation (RMSF <1.5 Å) values (Fig. 6B) demonstrated comparable system stability to the RBV-HEV-ORF1 complex. Among these compounds, Quercitrin, a flavonoid derivative, exhibited remarkable stability (RMSD: 1.93 Å), a finding consistent with previous reports of the high affinity between flavonoid and proteases in other hepatitis virus (Bachmetov et al., 2012; Shakya, 2019; Ugbaja et al., 2025). Structural stability metrics including Radius of Gyration (Rg) and Solvent Accessible Surface Area (SASA) analyses further confirm the maintenance of protein compactness during simulations, reinforcing the strength of these molecular interactions between HEV-ORF1 and the two hit ligands. These comprehensive computational results substantiate the potential of these flavonoid compounds as promising anti-HEV therapeutic candidates. These findings also align with previous research demonstrating the successful application of molecular docking and dynamics simulations approaches for identifying antiviral agents against HEV (Galani et al., 2021; Uddin et al., 2024).

In cell-based assays using the highly permissive Huh7.5 cell line, KHE demonstrated superior efficacy to ribavirin (RBV) in suppressing HEV replication, with an IC_50_ of 12.75 µg/mL compared to RBV’s 17.96 µg/mL (Fig. 7A). This finding aligns with previous reports on the antiviral activity of *K. grandifoliola* extracts and compounds. For instance, studies using HCV subgenomic replicons, pseudotyped particles, and cell-culture-derived HCV (HCVcc) revealed that *K. grandifoliola*’s methylene chloride/methanol extract and its fractions exhibit potent anti-HCV effects (Galani et al., 2016). Further investigation identified limonoids as key bioactive constituents that inhibit HCV replication via multiple mechanisms, including blockade of the CD81 receptor, downregulation of HCV-NS5B polymerase expression, and induction of the antiviral protein 2’,5’-oligoadenylate synthase-3 (OAS-3) (Kouam et al., 2021). Interestingly, KHE showed no cytotoxicity at concentrations ≤100 µg/mL, contrasting with RBV’s significant reduction in cell viability at 100 µg/mL (Fig. 7B). This favorable safety profile mirrors that of other plant-derived antivirals, (Lin et al., 2014), underscoring KHE’s potential therapeutic advantage. Our findings therefore suggest that KHE is likely non-toxic for the host cells and its anti-HEV activity may stem from the inhibition of the HEV-PLpro proteolytic activity, and potential induction of antiviral proteins such OAS-3. Collectively, our data position KHE as a promising source of active ingredients for developing broad-spectrum antiviral agents against viral hepatitis, including HEV. Future studies should therefore isolate and validate the individual contributions of top-scoring compounds, (Quercetin 3-[rhamnosyl-(1->2)-alpha-L-arabino-pyranoside] and Quercitrin), evaluate their *in vivo* efficacy and pharmacokinetics in humanized mouse model for HEV infection, and explore structure-activity relationships to optimize the top-scoring compounds for clinical translation.

## 5. Conclusion

This work elucidates the multi-faceted anti-HEV mechanism of KHE, from protease inhibition to viral replication suppression, driven by its diverse phytochemical arsenal. The superior safety and efficacy of KHE over RBV, combined with computational validation of key ligand-target interactions, highlights its potential as a promising source of antiviral agents against HEV, underlying the growing recognition of plant-derived compounds in antiviral therapy. By bridging gaps between traditional knowledge and modern pharmacology, this study contributes to the valorization of *K. grandifoliola* for the treatment of liver-related diseases and highlights its potential as reliable source of bioactive phytochemicals for combating viral hepatitis.

## Abbreviations

HCV: Hepatitis C virus HEV: Hepatitis E virus
HEV-PLpro: HEV papain-like cysteine protease
IPTG: Isopropyl-D-thiogalactoside
KHE: Hydro-ethanolic extract of *Khaya grandifoliola*
MDS: Molecular Dynamics Simulation
MTT: 3-(4,5-dimethylthiazol-2-yl)-2,5-diphenyl-2H-tetrazolium bromide
ORF: Open Reading Frame
RBV: Ribavirin
Rg: Radius of Gyration
RMSD: Root-Mean-Square Deviation
RMSF: Root-Mean-Square Fluctuation
SASA: Solvent-Accessible Surface Area
UHPLC-MS: Ultra high-performance liquid chromatography – Mass spectrometry

## Supporting information

Supplementary file S1: UHPLC-MS data file of the hydro-ethanolic extract of K. grandifoli

Supplementary file S2: Molecular dynamic simulations data file of the top-scoring complexes

## Acknowledgment

The authors are grateful to the Institute of Microbiology of the Chinese Academy of Sciences for providing the plasmids and the necessary facilities to conduct this research work.

## Availability of data and materials

All data generated or analyzed during this study are included in this published article and its supplementary information files.

## CRediT Authorship Contribution Statement

**Arnaud Fondjo Kouam:** Conceptualization, Investigation, Methodology, Software, Resources, Formal analysis, Validation, Supervision, Visualization, Project administration, Writing – original draft, Writing – review & editing

**Cromwel Tepap Zemnou:** Investigation, Methodology, Software, Formal analysis, Writing – original draft, Writing – review & editing.

**Armel Jackson Seukep:** Investigation, Methodology, supervision, validation, resources, Visualization, Writing – original draft, Writing – review & editing.

**Babayemi Olawale Oladejo:** Investigation, Methodology, Formal analysis, Resources, Visualization, Writing – review & editing

**Brice Fredy Nemg Simo:** Investigation, Methodology, Formal analysis, Validation, Visualization, Writing – review & editing.

**Kerinyuy Juliene Kongnyuy:** Investigation, Methodology, Formal analysis, Writing – review & editing.

**Armelle Gaelle Kwesseu Fepa:** Investigation, Methodology, Formal analysis, Writing – review & editing.

**Elisabeth Menkem Zeuko’o:** Investigation, Methodology, Formal analysis, Validation, Visualization, Writing – review & editing.

**Frédéric Nico Njayou:** Conceptualization, supervision, validation, resources, Visualization, Project administration, Writing – review & editing.

**Paul Fewou Moundipa:** Conceptualization, supervision, validation, resources, Visualization, Project administration, Writing – review & editing.

All authors read and approved the final version of the submitted manuscript.

## Supplementary information

Supplementary file S1: UHPLC-MS data file of the hydro-ethanolic extract of *K. grandifoliola*. Supplementary file S2: Molecular dynamic simulations data file of the top-scoring complexes

---

**Table 2:** Summary of kinetics parameters of HEV-PLpro without or with of RBV or KHE.

| Tested inhibitors<br>(µg/mL) |  | Michaelis-Menten representation |  | Lineweaver-Burk representation |  | Mechanism<br>of Inhibition | Inhibition constant |  |
| --- | --- | --- | --- | --- | --- | --- | --- | --- |
|  |  | V<sub>max</sub><br>(AMC µmol.min<sup>-1</sup>) | K<sub>m</sub><br>(µM) | V<sub>max</sub><br>(AMC µmol.min<sup>-1</sup>) | K<sub>m</sub><br>(µM) |  | (K<sub>ia</sub>, µg/mL) | (K<sub>ic</sub>, µg/mL) |
| Uninhibited<br>reaction (I<sub>0</sub> = 0) |  | 2.39 ± 0.03 | 29.29 ± 2.23 | 2.79 ± 0.09 | 31.74 ± 2.11 | / | / | / |
| RBV | I<sub>1</sub> = 4.5 | 2.38 ± 0.04 | 42.47 ± 1.85 | 2.79 ± 0.03 | 42.46 ± 2.16 | Competitive<br>inhibition | 11.63 ± 1.40 | / |
|  | I<sub>2</sub> = 9 | 2.38 ± 0.02 | 55.65 ± 2.08 | 2.79 ± 0.04 | 55.64 ± 1.35 |  |  |  |
| KHE | I<sub>1</sub> = 5 | 2.16 ± 0.03 | 37.85 ± 1.70 | 2.35 ± 0.02 | 38.19 ± 2.15 | Mixed<br>inhibition | 31.28 ± 0.91 | 24.44 ± 1.61 |
|  | I<sub>2</sub> = 10 | 2.00 ± 0.02 | 42.52 ± 2.13 | 2.06 ± 0.04 | 43.61 ± 1.27 |  |  |  |

Data are means ± SD of three independent experiments in triplicate; **RBV**: Ribavirin; **KHE**: Hydro-ethanolic extract of *K. grandifoliola*; **V<sub>max</sub>**: Maximum reaction velocity; **K<sub>m</sub>**: Michaelis-Menten constant; **K<sub>ia</sub>**: Inhibition constant toward K<sub>m</sub>; **K<sub>ic</sub>**: Inhibition constant toward V<sub>max</sub>.

**Table 3:** Molecular Docking scores and interactions between amino acids residues of HEV_OFR1 and the best ligands identified from KHE.

| HEV_ORF1 (PDB ID: 6NU9) |  |  |  |  |  |  |  |
| --- | --- | --- | --- | --- | --- | --- | --- |
| Ligands | Docking scores (Kcal/mol) | Hydrogen Bonds with specific amino acid residues |  |  | Hydrophobic interactions with specific amino acid residues |  |  |
|  |  | Conventional | Pi-Donor / Acceptor /Unfavorable | Carbon | Alkyl/Amide-Pi/Pipi-T | Pi-Alkyl | Pi-Sigma/Anion/Lone |
| L3 | -7.07 | 1 with Lys63 | 0 | 1 with Tyr146<br>1 with Val180<br>1 with Glu182 | 2 with His164 | 1 with Phe65 | 1 with His164 |
|  |  | 1 with Thr64 |  |  |  | 1 with His164 |  |
|  |  | 1 with Asn149 |  |  |  | 2 with Pro165 |  |
|  |  | 2 with Val180 |  |  |  | 1 with Val180 |  |
|  |  | 1 with Glu182 |  |  |  |  |  |
| L4 | -6.23 | 1 with Ser68 | 1 with His164 | 1 with Gly178 | 0 | 1 with Pro177 | 2 with Arg66 |
|  |  | 2 with Glu166 |  |  |  |  |  |
| L10 | -7.68 | 1 with His164 | 1 with Pro177 | 1 with Asp55 | 0 | 1 with Ala73<br>1 with Pro177 | 1 with Arg66 |
|  |  | 1 with Glu166 |  | 1 with Ser68 |  |  |  |
|  |  | 2 with Glu182 |  | 1 with Gly178 |  |  |  |
|  |  |  |  | 1 with Glu182 |  |  |  |
| L12 | -7.43 | 2 with Arg66 | 0 | 1 with Arg66<br>1 with Glu182 | 0 | 1 with Pro177 | 1 with Ser68 |
|  |  | 1 with Ser68 |  |  |  |  |  |
|  |  | 2 with Glu166 |  |  |  |  |  |
|  |  | 1 with Glu182 |  |  |  |  |  |
| RBV | -4.87 | 1 with Asp55 | 0 | 1 with Ser68 | 1 with Thr67 | 1 with Arg66 | 0 |
|  |  | 2 with Arg66 |  | 1 with His179 |  |  |  |
|  |  | 1 with Thr67 |  | 1 with Trp181 |  |  |  |
|  |  | 1 with Trp181 |  |  |  |  |  |

L3: Scopolin; L4: Procyanidin (catechin+epicatechin); L10: Quercetin 3-[rhamnosyl-(1->2)-alpha-L-arabinopyranoside]; L12: Quercitrin. RBV: Ribavirin

